# Homologous BHLH transcription factors induce distinct deformations of torsionally-stressed DNA: a potential transcription regulation mechanism

**DOI:** 10.1101/2022.01.07.475332

**Authors:** Johanna Hörberg, Kevin Moreau, Anna Reymer

## Abstract

Changing torsional restraints on DNA is essential for the regulation of transcription. Torsional stress, introduced by RNA polymerase, can propagate along chromatin facilitating topological transitions and modulating the specific binding of transcription factors (TFs) to DNA. Despite the importance, the mechanistic details on how torsional stress impacts the TFs-DNA complexation remain scarce. Herein we address the impact of torsional stress on DNA complexation with homologous human basic-helix-loop-helix (BHLH) hetero- and homodimers: MycMax, MadMax, and MaxMax. The three TF dimers exhibit specificity towards the same DNA consensus sequences, the E-box response element, while regulating different transcriptional pathways. Using microseconds-long atomistic molecular dynamics simulations together with the torsional restraint that controls DNA total helical twist, we gradually over- and underwind naked and complexed DNA to a maximum of ±5°/b.p. step. We observe that the binding of the BHLH dimers results in a similar increase in DNA torsional rigidity. However, under torsional stress the BHLH dimers induce distinct DNA deformations, characterised by changes in DNA grooves geometry and a significant asymmetric DNA bending. Supported by bioinformatics analyses, our data suggest that torsional stress may contribute to the execution of differential transcriptional programs of the homologous TFs by modulating their collaborative interactions.

## Introduction

Changing torsional constraints on DNA comprise one of the major regulatory forces of eukaryotic transcriptional control.(Lavelle 2008; Ma, Bai, and Wang 2013; Corless and Gilbert 2016; Kouzine et al. 2013; Naughton et al. 2013; Fogg et al. 2021) In transcription, torsional strain is primarily introduced by RNA polymerase which forces DNA to rotate around its axis as the molecule threads through the transcription machinery.(Osborne et al. 2004; Boeger et al. 2005) The imposed torsional strain underwinds and overwinds DNA upstream and downstream of a transcribed gene, correspondingly, and can propagate along the chromatin fibre. The speeds and ranges of torsional strain propagation depend on the underlying nucleotide sequence.(Naughton et al. 2013) When propagating along the chromatin fibre torsional stain brings changes to the genome organization: locally introducing DNA bending and twisting, which affects the stability of nucleosome core particles,(Teves and Henikoff 2014) and higher-order chromatin structures.(Corless and Gilbert 2016) Through these changes, torsional strain can modulate transcription of near-located genes. Generally, genes that experience torsional strain are more efficiently transcribed.(Weintraub, Cheng, and Conrad 1986; Dunaway and Ostrander 1993) Furthermore, the torsional strain might be necessary for the initiation of transcription,(Dunaway and Ostrander 1993; Mizutani et al. 1991; Schultz et al. 1992; Tabuchi and Hirose 1988; Mizutani, Ura, and Hirose 1991) as DNA underwinding appears essential for the melting of the TATA-box sequence near transcription start sites.(Liebl and Zacharias 2020) To complete the picture of how torsional strain contributes to eukaryotic transcriptional control, we must also understand how torsional strain impacts DNA specific binding by transcription factor proteins.(Noy, Sutthibutpong, and A. Harris 2016; Pyne et al. 2021; J. Hörberg and Reymer 2020)

Locally DNA responds to torsional strain in a highly sequence-specific manner. Certain dinucleotide steps, depending on their flanking environment,(Liebl and Zacharias 2020; Kannan, Kohlhoff, and Zacharias 2006; Liebl and Zacharias 2017; Reymer, Zakrzewska, and Lavery 2018; Johanna Hörberg and Reymer 2018) can effectively absorb both negative and positive torsional strain by switching their twist. These conformational transitions allow the rest of the DNA molecule to preserve a B-like conformation. The “twist-capacitors” dinucleotides, which include pyrimidine-purine (YpR) and purine-purine (RpR) steps, appear to regulate protein-DNA complexation, as twisting transitions are coupled to changes in shift and slide(Dans et al. 2012; Pasi et al. 2014; Dans et al. 2019; Balaceanu et al. 2019)– helical parameters important for the protein-DNA readout mechanism.(Johanna Hörberg et al. 2021) Previously we addressed the impact of torsional strain on DNA complexation with a human basic-leucine-zipper (BZIP) transcription factor (TF) MafB.(J. Hörberg and Reymer 2020) When specifically bound to its DNA target, the protein locks the twist-capacitor dinucleotides in one conformational sub-state, favourable for the complexation. Consequently, the energy cost for DNA twisting almost doubles – suggesting that BZIP factors may hinder the propagation of torsional strain along DNA, potentially regulating the cooperative binding of collaborative TFs, or contributing to alterations in genome topology. However, these results provide only first insights; a complete understanding of the mechanistic aspects of how torsional strain affects TF-DNA complexation remains limited. TF-DNA complexes involving other families of TFs may respond differently to torsional stress. Also, there might be differences among homologous TFs.

We herein address the impact of torsional strain on DNA complexation with three homologous basic helix-loop-helix (BHLH) TFs dimers, MycMax, MadMax, and MaxMax. The BHLH factors is one of the most abundant families of eucaryotic TFs that modulate cell proliferation, differentiation, and apoptosis.(Atchley and Fitch 1997; Dennis, Han, and Schuurmans 2019; De Masi et al. 2011; de Martin, Sodaei, and Santpere 2021) The MycMax, MadMax, and MaxMax dimers exhibit specificity towards identical DNA consensus sequences – yet regulate different transcriptional pathways.(Grandori et al. 2000; Diolaiti et al. 2015; Giardino Torchia and Ashwell 2018) We study MaxMax, MycMax, and MadMax dimers bound to the palindromic sequence ‘5’-GGCGAGTAG**CACGTG**CTACTCGC-3’, containing the E-box response element (in bold) under relaxed and torsionally constrained conditions using microsecond-long umbrella sampling molecular dynamics (MD) simulations. With the torsional restraint that controls the total twist of DNA, without restricting other degrees of freedom, we gradually over- and underwind DNA to a maximum of ±5°/base pair (b.p.) step. We observe that, relative to unbound DNA, the BHLH factors make DNA more torsionally rigid, resulting in similar torsional moduli for complexed DNA. However, under torsional stress, the BHLH factors induce distinct DNA deformations. We complement our mechanistic studies with bioinformatics analysis. Using CHIP-seq data on several cell lines, we analyse the binding properties of Myc, Max, and Mad proteins, investigating their relative position with respect to the nearest nucleosomes and transcription starting sites, as well co-binding with other TFs. Our results suggest: distinct responses to DNA twisting by homologous TFs may be a complementary mechanism to variations in transactivation domains (TAD), contributing to the recruitment of different collaborative TFs and, subsequently, differential regulatory responses.

## Methods

### Simulated Systems

Four systems: MycMax- (PDB ID: 1NKP)(Nair and Burley 2003), MadMax- (PDB ID: 1NLW)(Nair and Burley 2003), MaxMax-bound DNA (PDB ID: 1AN2)(Ferré-D’Amaré et al. 1993) and free DNA in B-form were studied. All systems contain a DNA 23-mer: GGCGAGTAG**CACGTG**CTACTCGC, with the E-box region in bold. USCF Chimera(Pettersen et al. 2004) was used to modify the DNA sequence of the MaxMax-DNA complex to match that of the MycMax-DNA and MadMax-DNA complexes; and to determine the protonation state of His residues. JUMNA(Lavery, Zakrzewska, and Sklenar 1995) program was then used to extend the flanking sites and to relax bad protein-DNA contacts.

### Molecular Dynamics Simulation Protocol

All molecular dynamics (MD) simulations are performed with the MD engine GROMACS v2019.4(Abraham et al. 2015) using same protocol as described previously.(J. Hörberg and Reymer 2020) In the restrained MD simulations of cascade umbrella sampling, we use the inhouse developed torsional restraint(Reymer, Zakrzewska, and Lavery 2018) that controls the total twist of a DNA fragment. The restrained sets the desired value of *twist_ref_* using a simple quadratic function, *K_tw_ (twist-twist_ref_)^2^*, implemented via PLUMED v2.5.3.(Bonomi et al. 2009) We use the force constant (*K_tw_*) of 0.06 kcal mol^-1^ degrees^-2^, the smallest value that provides the desired twist without inducing any structural artefacts. In all simulations we use AMBER 14SB(Maier et al. 2015) and Parmbsc1(Ivani et al. 2016) force fields to treat the protein and DNA, respectively.

The protein-DNA complexes and free DNA oligomer are separately solvated in cubic periodic boxes by SPC/E(Mark and Nilsson 2001) water molecules with a buffer distance of 12 Å to the walls. The systems are first neutralized by K+ counterions, then additional K+ and Cl- ions are added to reach a physiological salt-concentration of 150 mM. Applying periodic boundary conditions, each system is subjected to energy minimization with 5000 steps of steepest descent, followed by 500 ps equilibration-runs with week position restraints on heavy solute atoms (1000 kJ/mol) in the NVT and NPT ensembles to adjust temperature(Berendsen et al. 1984) and pressure(Parrinello and Rahman 1981) to 300 K and 1 atm. Releasing the restraints, 0.6 μs simulations are then carried out at constant pressure and temperature (1 atm and 300 K).

Following the unrestrained MD simulations, the cascade umbrella sampling(Torrie and Valleau 1977) is performed with 0.5 μs sampling time per umbrella window to allow sufficient convergence of DNA conformational substates and ion populations. We apply the twist restraint to the central E-box region and the four adjacent 5’- and 3’-flanking nucleotides; 13 b.p. steps in total. Starting from a relaxed state, the total twist of the restrained fragment is gradually changed by ±0.5°/b.p step (±6.5° in total per umbrella window), until a maximum overwound and underwound state of 5°/b.p. step is reached. The final structure from every window is used as the starting point for the following umbrella window. The Weighted Histogram Analysis Method (WHAM)(Kumar et al. 1992), implemented in PLUMED is used to derive the potential of mean force (PMF) with respect to DNA twisting. The total simulation time for each system is 11.1 mks.

### Elastic Force Constant Analysis

Quadratic regression analysis in MatLab is used to obtain the twisting force constants, K (kcal/mol deg^2^) from the PMF profiles. The analysis is performed for the regions corresponding to a Δtw of ±2°, and for over- and undertwisting individually to highlight any asymmetry of the PMF profiles. The derived force constants are used to calculate DNA torsional modulus according to the isotropic rod model, given by equation (1); where T is the torque that results from a change in twist Δ***θ*** over a standard b.p. length L (0.34 nm). From the torsional modulus we derive the torsional persistence length P using equation (2), using k_B_T at room temperature.

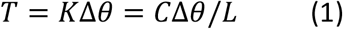

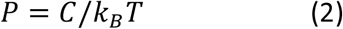

### Trajectory analysis

Curves+ and Canal(Lavery et al. 2009) programs are used to derive DNA helical parameters, backbone torsional angles and groove geometry parameters for each trajectory snapshot extracted at 1 ps intervals. A multivariate Ising model is used to calculate DNA deformation energies. The CPPTRAJ(Roe and Cheatham 2013) tool from AMBERTOOLS16 software package is used to derive specific and nonspecific protein-DNA contacts along the different torsional regimes as described previously.(J. Hörberg and Reymer 2020)

### Bioinformatic Analysis

Analysis of the nearest binding partners of Myc and Max was performed using TFregulomeR(Lin et al. 2020) package on ChIP-seq datasets from multiple cell lines (A549, NB4, K562) downloaded from TFregulomeR data compendium. The output gives, among other information, the top-10 co-binders (ChIP-seq signal around +/-100bp of the peaks summits, corresponding the to the binding of the studied TFs). The TFregulomeR package allows to differentiate the Myc and Max binding distribution corresponding district genomic regions such as promoter, introns, exons, intergenic regions, etc.

### Additional Information

MatLab and R software(Core Team 2013) were used for post-processing and plotting of all data. USCF Chimera(Pettersen et al. 2004) was used for creating molecular graphics.

## Results

### Torsional Moduli of Free and Complexed DNA

To address how changing torsional restraints on DNA impact the molecular complexation with homologous transcription factors and consequently their transcription regulatory programs, we select homo- and heterodimers of the Myc/Max/Mad network. The Myc/Max/Mad proteins belong to the basic helix-loop-helix (BHLH) family of eukaryotic transcription factors (TFs) that exhibit specificity towards same DNA response elements but paly distinct roles in transcriptional control. Upon association with its DNA target sites, the MycMax dimer acts as a transcriptional activator, which induces histone acetylation. The MadMax dimer antagonizes MycMax and acts as a transcriptional repressor, which recruits histone deacetylases. The MaxMax dimer also antagonizes MycMax, however, since Max lacks a transactivation/transrepression domain, it is considered transcriptionally inert.(Grandori et al. 2000)

We first subject the MaxMax-, MycMax-, and MadMax-DNA complexes and free DNA, containing the E-box response element sequence (‘GGCGAGTAG**CACGTG**CTACTCGC’) (Figure S1) to unrestrained MD simulations. We continue with cascade umbrella sampling using the torsional restraint that controls total twist of a restrained DNA region. The restraint is applied to the E-Box response element and the four adjacent 5’- and 3’ flanking nucleotides, 13 b.p. steps in total. Starting from the relaxed duplex, we gradually over- and underwind free and protein-bound DNA until reaching a maximum of ±5°/b.p. step (corresponding to ± 0.15/b.p. step in supercoiling density). In accordance with previous studies, at all simulated torsional regimes the restrained DNA region in the four systems preserves a B-like conformation. We observe no significant DNA b.p. flipping or melting, as supported by DNA stretch and opening distributions (Figures S5A-D and S6A-D). The selected torsional restraints range, though potentially exaggerated, represents an approximation of extreme local changes that may arise near transcription starting sites upon transcription initiation; and allows us to gain mechanistic insights into these highly dynamic aspects of eukaryotic transcriptional control. The ultimate range of *in vivo* changes in torsional restraints on DNA remains to be determined.

From the torsionally restrained MD simulations we derive the potential of mean force (PMF) profiles showing the free energy cost for DNA twisting transitions (Figure 1A, Figure S3). To compare the PMF profiles, we plot the changes in average twist per b.p. step with respect to the relaxed average twist that varies from 34.0 for MadMax-bound DNA to 34.7 for MycMax-bound DNA (Table S1). To compare the torsional rigidity of DNA in the four systems, we derive the torsional force constants, torsional moduli, and persistence lengths using quadratic regression (Figure S7, Table S1). We observe that the binding of the BHLH dimers make DNA more torsionally rigid, resulting in the torsional moduli of 628, 691, and 672 pN nm^2^ for MycMax-, MadMax-, and MaxMax-bound DNA, correspondingly, vs 457 pN nm^2^ for free DNA. The observation is consistent with our previous study of MafB,(J. Hörberg and Reymer 2020) where we showed that the TF binding almost doubles the torsional rigidity of DNA (853 pN nm^2^ for complexed vs 442 pN nm^2^ for free DNA), reflecting the differences in the complexation mechanisms between the BZIP and BHLH families of transcription factors.

**Figure 1:**
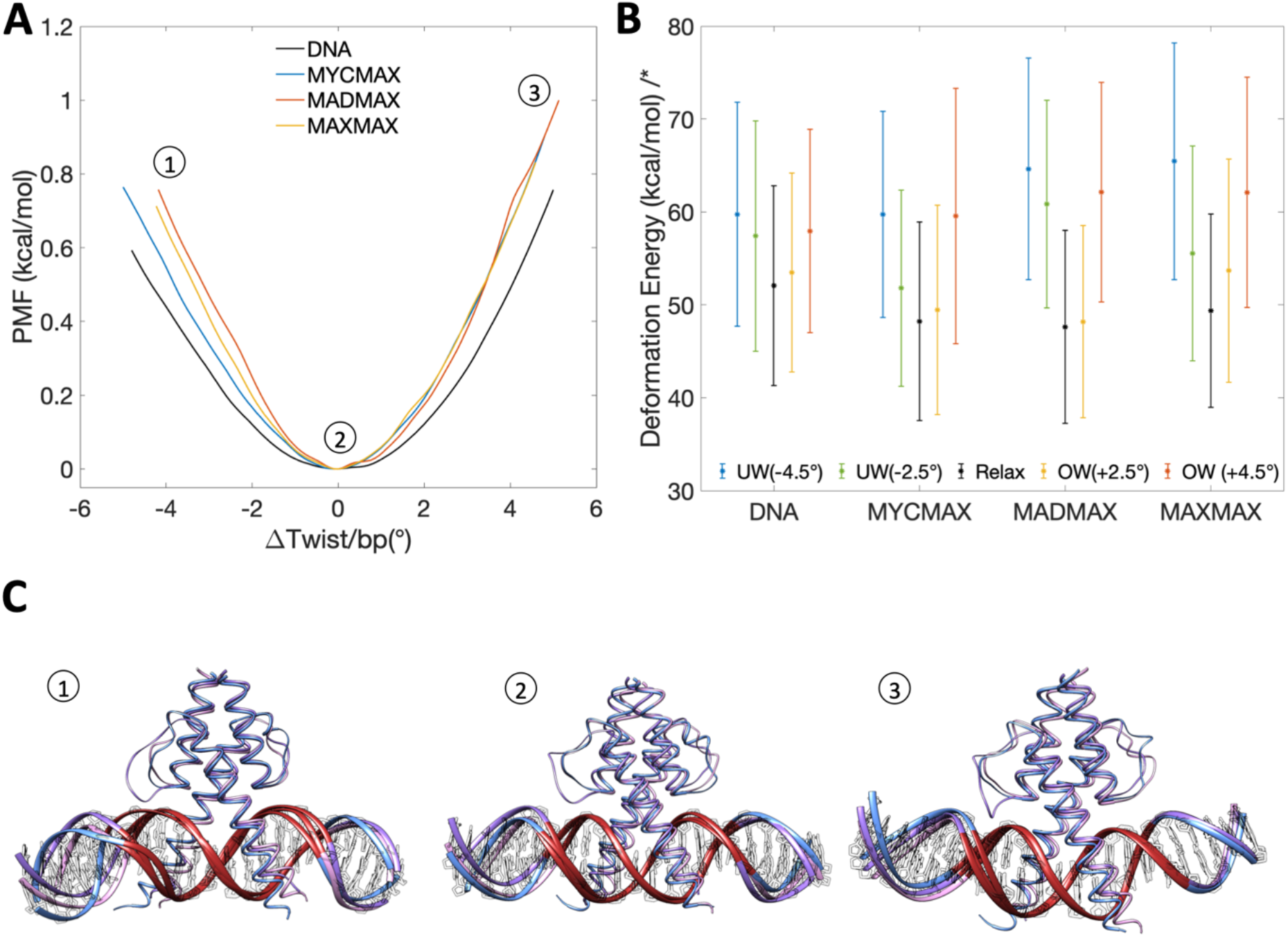
**A:** PMF profiles showing the energy cost for twisting transitions for free and BHLH-bound DNA. **B:** MycMax-, MadMax- and MaxMax induced DNA deformations are shown for (1) underwound regime, (2) torsionally relaxed regime and (3) overwound regime; with the restrained DNA region in red, MycMax in blue, MadMax in magenta, and MaxMax in pink. **C:** DNA deformation energy of the restrained region (GTAG**CACGTG**CTAC, E-box response element in bold) calculated with a multivariate Ising mode.(Liebl and Zacharias 2021)

Despite the similar increased DNA torsional rigidity, induced by the protein complexation, we observe that the three BHLH dimers deform DNA in a different fashion. Starting from −1.0°/b.p step during the underwinding regime, we observe that DNA molecules experience a local bending induced by an increased roll angle at several b.p. steps and changed geometry of DNA grooves (Figures 1B and S3, Videos S1-4). The torsionally induced deformations differ also between free and protein-bound DNA (Figure S3). For free DNA the imposed torsional stress is evenly distributed over the entire restrained region (Figure 2), with no significant bending even at higher torque regimes (Figure S4). While for protein-bound DNA (see details below), the imposed torsional stress is mostly accumulated in the flanking regions outside the E-box response element (Figure 2), where the observed deformations are predominantly localised.

**Figure 2:**
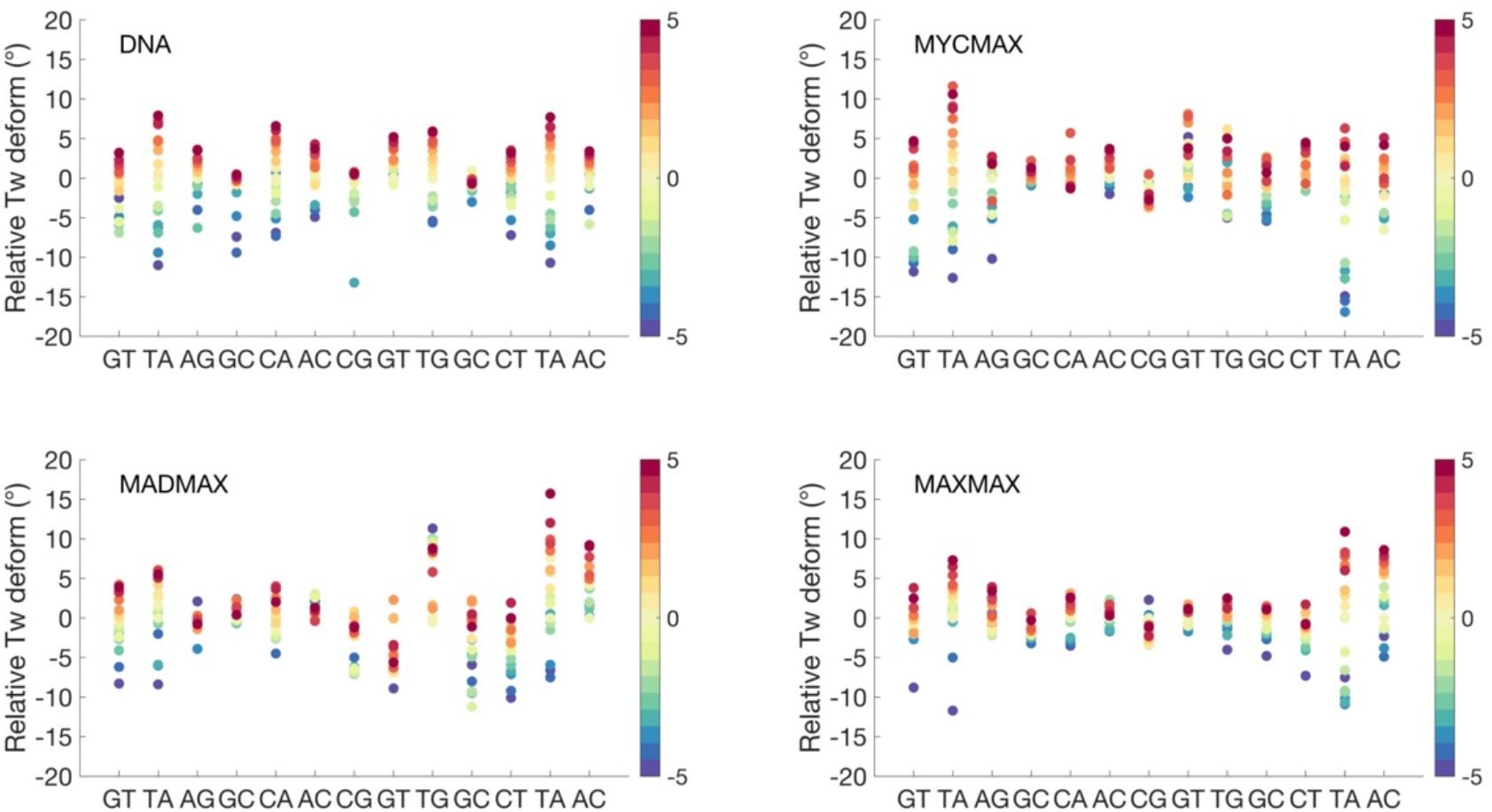
Changes of twist angles for the restrained DNA-region in unbound and MycMax-, MadMax- and MaxMax-bound DNA as a function of the requested average change of twist per base pair step, indicated by a colourbar to the right.

We also calculate the deformation energies (Figure 1C) for the restrained DNA regions using a multivariate Ising model.(Liebl and Zacharias 2021) Consequently, in all four systems the deformation energies increase as the torsional restraints are applied, both for under- and overtwisting. However, the increase in deformation energies is system-specific, suggesting that torsional stress may modulate the differential transcriptional response of homologous BHLH TFs by differently impacting the local conformational flexibility of DNA.

### Torsional Stress-Induced Changes in DNA Structure

To understand the BHLH dimer specific torque-induced DNA deformations, we first analyse the contributions of individual b.p. steps to the absorption of the imposed torsional stress (Figures 2 and S8). To avoid any boundary effects, we exclude from the analysis the outer b.p. steps (GpT and ApC) of the restrained region (GTAG**CACGTG**CTAC, the E-box response element in bold). Contrary to free DNA, where the imposed torsional stress is absorbed by the flexible pyrimidine-purine steps, TpA and CpA, of the palindrome (G**TA**G**CA**CG**TG**C**TA**C), for protein-bound DNA the CpA and TpG steps within the E-box sequence become torsionally-rigid. As a result, the torque accumulates mostly in the flanking regions. For MycMax- and MaxMax-bound DNA, the flanking TpA steps dominate the absorption of both negative and positive torque (Figure 2). However, as the TpA steps exhibit a preference for a high twist state under the relaxed conditions (Figure S8), they are less efficient in absorbing positive torsional stress, contributing to an increased energy cost for DNA overtwisting. For MadMax-bound DNA the first flanking TpA step remains rigid, while the second flanking TpA step efficiently absorbs only positive torsional stress as it prefers a low twist state under the relaxed conditions (Figures 2 and S8). Thus, the imposed negative torsional stress is distributed among the less torsionally flexible b.p. steps in the second halfsite of the restrained DNA region, which increases the energy cost for DNA undertwisting.

For the torsionally active b.p. steps, behaviour of other translational and rotational b.p. steps parameters under torsional stress follow the earlier reports: torsional stress brings changes in roll, shift, and slide – which are coupled to twist via BI/BII backbone conformational transitions (Figures S9-15). Roll shows negative correlation to twist, and slide – positive. Behaviour of the shift parameter appears more complexed. In the relaxed state, we observe shift bimodality/multimodality for the torsionally flexible b.p. steps, which changes as DNA undergoes under- and overtwisting.

We further characterise the differences in the BHLH dimer-specific torque-induced DNA deformations, by analysing DNA axes bending (Figures 3 and S4) and groove parameters (Figure 4). For all protein bound DNA we observe an asymmetric bending towards the major groove during the underwinding regime. The axis bending for MycMax- and MaxMax-bound DNA, from the Myc- and Max_1-side, correspondingly, gradually increases with undertwisting up to 5°per b.p. step (Figures 3, S4 and Videos S2-S3). Max_1 of the MaxMax homodimer refers to the monomer, which forms a greater number of specific contacts with DNA (see Protein-DNA contacts for further details). Contrary, for MadMax-bound DNA, similarly to free DNA, we observe no significant DNA bending during underwinding. Overwinding also contributes to an increase in axis bending for all BHLH-bound and free DNA. However, the changes are smaller, about 2°per b.p. step, and more uniformly distributed over the restrained region, resulting in a symmetric smooth bending towards the minor groove (Figure S3, Video S2-S3). For DNA groove parameters, DNA underwinding results in an increase in the major groove width and depth, and a decrease in the minor groove depth. The reverse occurs during overwinding. For BHLH-bound DNA, the changes in the grooves geometry are more significant at the flanking sites that accumulate more of the imposed torsional stress, Myc-, Max_1-, and Mad-side, correspondingly.

**Figure 3:**
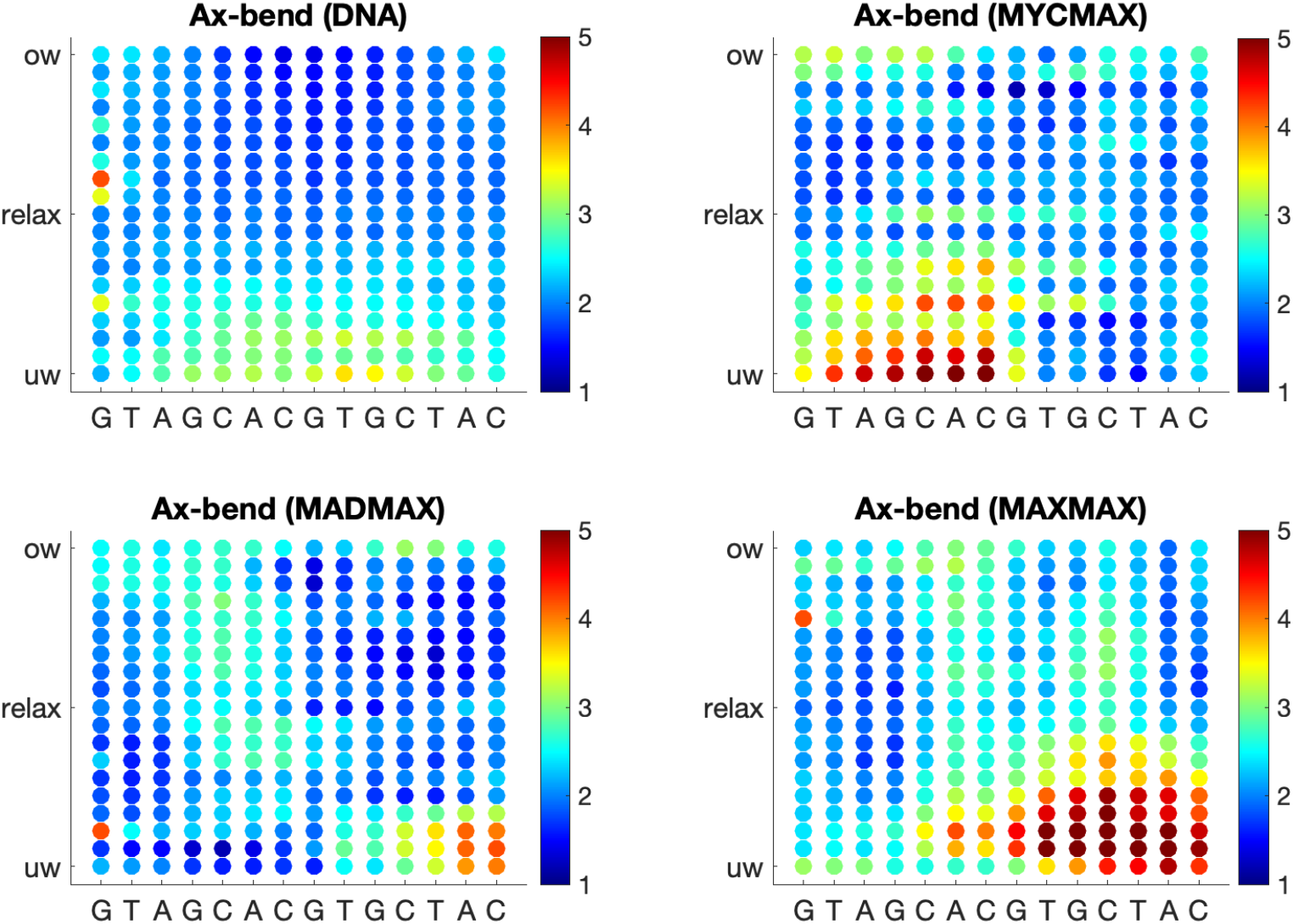
Change in average ax-bending of b.p. of the restrained DNA region (GTAGCACGTGCTAC) along the torsional regimes denoted with a colourbar.

**Figure 4:**
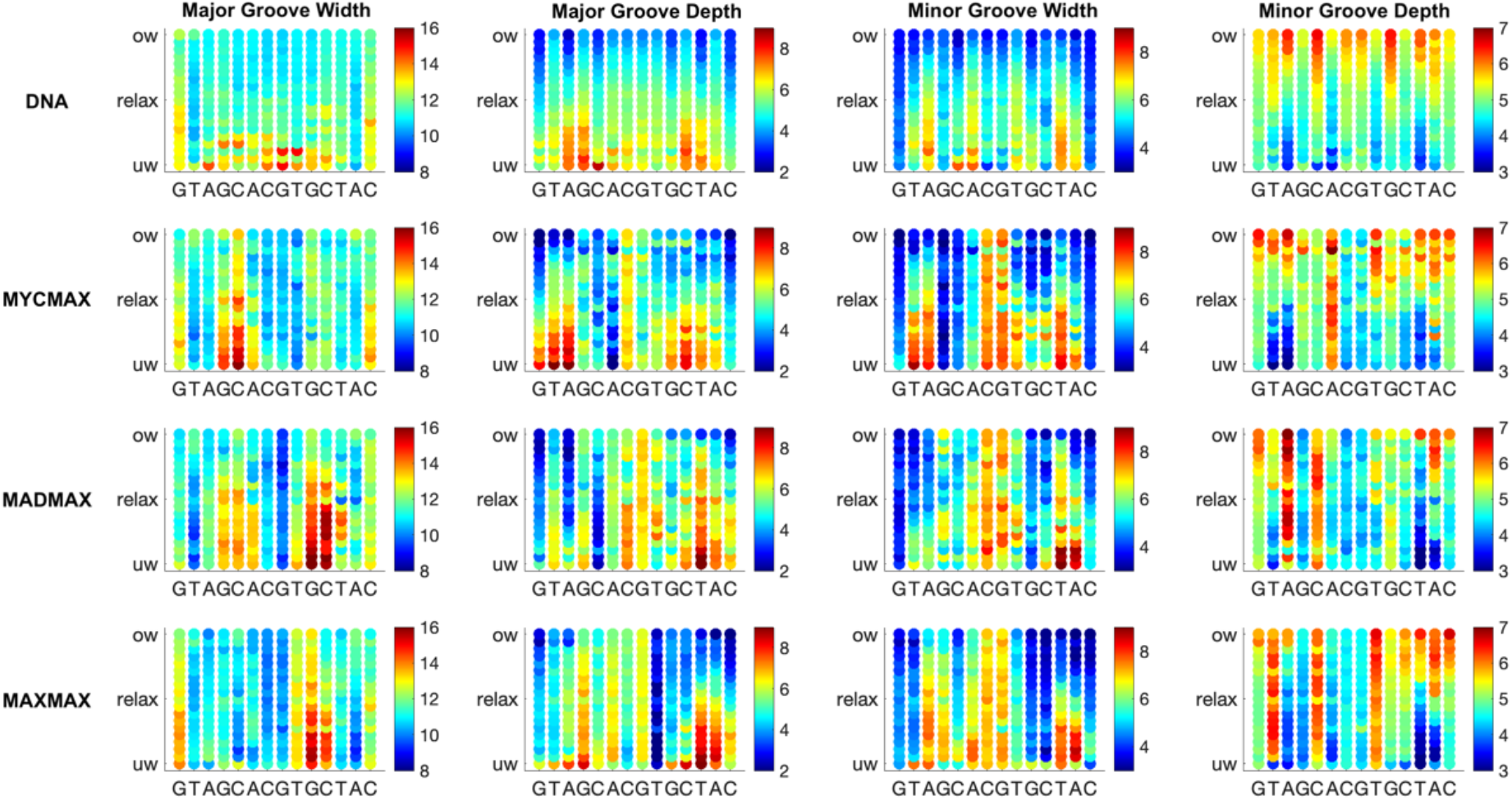
Change in average groove parameters of the restrained DNA region (GTAGCACGTGCTAC) along the torsional regimes denoted with a colourbar.

### Protein-DNA Contacts Networks at Different Torsional Stress Regimes

We continue with the analysis of differences in the intermolecular contacts networks exploited by the three BHLH dimers. To recognise their DNA targets, the BHLH family utilizes a five-residues motif (**\*\***xx**E**xx**R\***)(De Masi et al. 2011). For Myc, Mad, and Max the five-residues motif corresponds to **HN**xx**E**xx**RR** (Figure S1). Upon DNA binding, one of the monomers of the BHLH dimers forms further specific contacts with the E-box sequence, this includes Myc of MycMax, Mad of MadMax, and Max_1 monomer of MaxMax (Figures 5, S17A, S19). The other monomer interacts with DNA more nonspecifically. The Myc/Mad/Max_1 monomer shows nearly identical networks of specific intermolecular contacts (Figures 3 and S19). In the torsionally relaxed state, from the five-residues motif (HNxxExxRR), histidine interacts specifically with the TG b.p. step on the opposite strand of the E-box half-site (CACG/CG**TG**); asparagine with the T base on the opposite strand of the E-box half-site (CACG/CG**T**G), glutamate with the CA b.p. step (**CA**CG/CGTG) and the T base on the opposite strand of the E-box half-site (CACG/CG**T**G); first arginine with the CA b.p. step (**CA**CG/CGTG) and the flanking sites; and second arginine with the central CG b.p. step on the opposite strand of the E-box half-site (GCACG/**CG**TG). Furthermore, the MadMax dimer has an additional specific contact, the Arg91 residue of the Mad loop interacts with the flanking TA step (G**TA**GCACG) from the minor groove (Figure S18-19B), which explains the torsional rigidity of the step.

**Figure 5:**
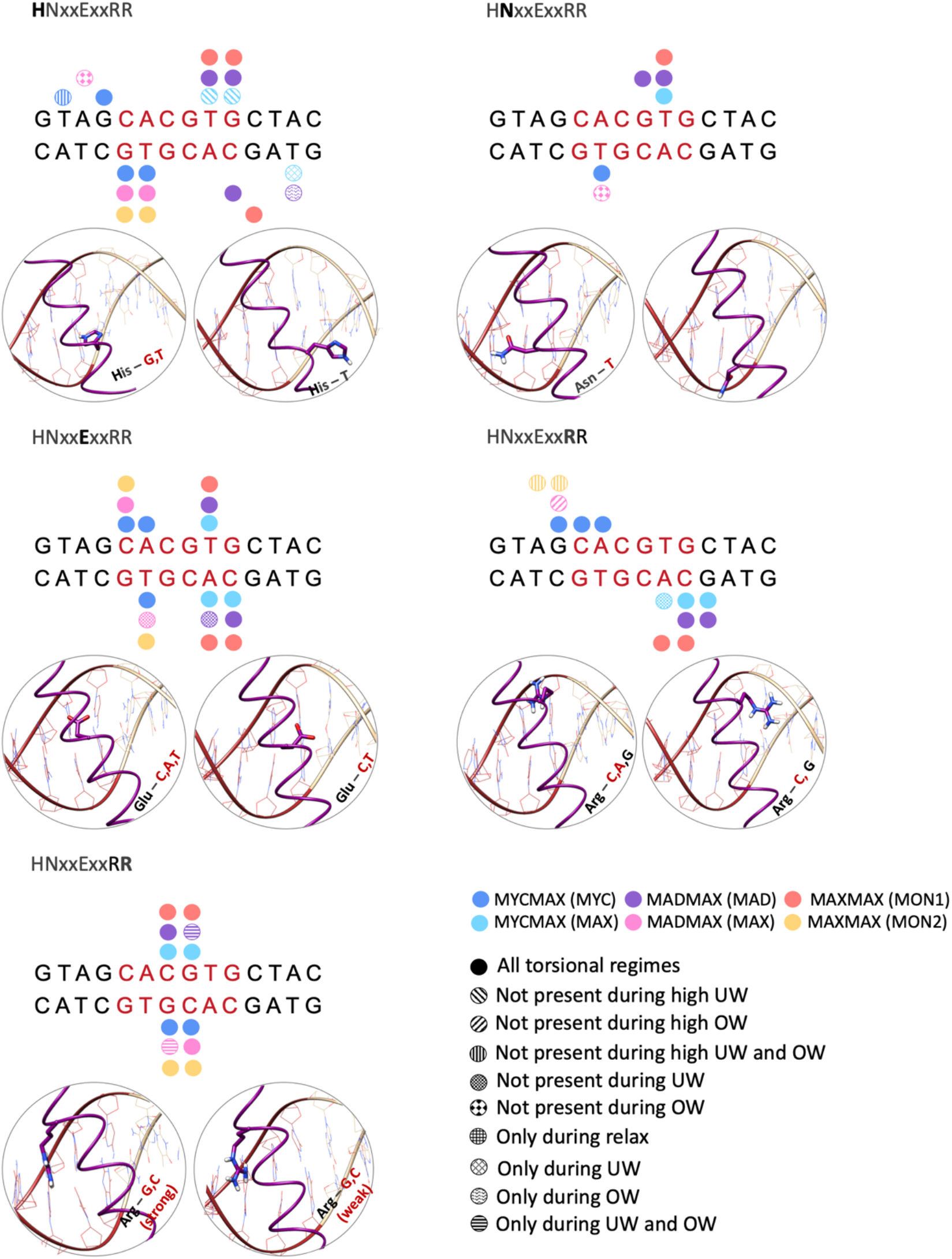
Specific contacts exploited by the five residues motif HNxxExxRR during the different torsional regimes.

The intermolecular specific interactions by the Max monomer, including Max of the MycMax and MadMax heterodimers, and Max_2 monomer of the Max-homodimer, follow to some extend the above-described specific contacts (Figures 3, S18, and S19). The differences include, for Max of MadMax and Max_2 of MaxMax, the Glu residue shows no interactions with the CA b.p. step; the first Arg residue interacts only with the flanking sites; for the MadMax dimer the second Arg residue oscillates between specific and nonspecific interactions with the CG b.p. step (Figure S18).

We further analyse the evolution of the strength of specific and nonspecific contacts, and the total number of protein-DNA contacts along the different torque regimes (Figures S17-18). The analyses show that most of the protein-DNA contacts remain stable over the different degrees of torsional strain. There are, however, BHLH dimer- and torque-specific differences in the protein-DNA contacts that can explain the observed differences in DNA torque-induced deformations during the underwinding regime. Upon high level of underwinding (> −3°/b.p.) the His residue (**H**NxxExxRR) of the Max monomer of the MycMax heterodimer rearranges its orientation to hydrophobically interact with the T base (step (GTAGCACGTGC**T**AC) Figure S19A), which limits the torsional activity of the corresponding TA step, leading to a smaller change in the roll angle and the major groove width. This in addition to a greater number of nonspecific interactions from the Max side creates potential steric clashes that hinders DNA bending. Similarly, during the underwinding regime the nonspecific interactions that involve the His27 and Leu31 residues of Max_2 in the MaxMax homodimer, and the Ser 55-57 residues of Mad in the MadMax heterodimer prevent DNA bending from the corresponding sites.

There are also torque-specific changes in the protein-DNA specific interactions that have no impact on the observed torque-induced BHLH dimer-specific DNA deformations. Those include the Glu-A specific contacts by Max of MycMax and Mad of MadMax that become weaker upon underwinding (Figure S18), the second Arg-G specific contact for Mad and Max in MadMax that becomes stronger upon underwinding.

We also observe fluctuations in the ratio of the specific to nonspecific contacts and their strengths, which are linked to the flickering power of the long side chain residues, oscillating between interactions with DNA bases and backbone. The overall stability of the intermolecular contact networks reflects that the BHLH factors form mainly contacts with the E-box response element, while the imposed torque is predominantly absorbed by the flanking regions.

## Discussion

Our results show that homologous BHLH dimers, MycMax, MadMax, and MaxMax can hinder the propagation of torsional stress along the genome by making the bound E-box sequence more torsionally rigid. Consequently, the imposed torque is accumulated at the flanking sites, resulting in the distinct BHLH dimer-specific DNA deformations during the underwinding regime. The negative torque-induced deformations, characterised by changes in DNA grooves geometry and an asymmetric bending of the E-box half side flanks, are more significant for MaxMax and MycMax DNA. The deformations occur at the Myc and Max_1 side, as these monomers form more base-specific contacts than their dimer partners, Max and Max_2, correspondingly. Despite experiencing the distinct deformations, the increase in DNA rigidity is relatively similar (see Table S1). Here, we want to point out that we study only one DNA sequence; it is likely that the observed DNA deformations are sequence specific. In addition, the studied DNA sequence is relatively short, for longer DNA the deformations may show a different amplitude as well as localisation.

Our results allow us to hypothesise that the torque-induced BHLH dimer-specific DNA deformations can contribute to the TFs differential transcriptional responses by producing binding sites for distinct collaborative proteins. To validate our hypothesis, we explore potential binding partners of the BHLH dimers, using the ChIP-seq data available for Myc and Max TFs. The analysis reveals that among ten most frequent binding partners, within a range of 200 b.p., there are members of E2F, BZIP, and Zinc finger TFs families (Figure S19A-B). Additionally, previous studies also listed the TATA-box binding protein(Wei et al. 2019; Lourenco et al. 2021), TBP, as a frequent co-binder of the MycMax dimer. Interestingly, the TBP and the E2F-factors deform DNA in a similar fashion as the one we observe for the restrained Max_1 and Myc-flanks of MaxMax- and MycMax-DNA, correspondingly, during the underwinding regime. Thus, potentially the binding of the MycMax/MaxMax dimers could facilitate the binding of the TBP and E2F factors. For BZIPs and Zinc fingers, which induce no major conformational change of DNA but due to the flexibility of their DNA binding domains may associate with significantly deformed DNA(J. Hörberg and Reymer 2020; Patel et al. 2018), the co-binding with MycMax/MaxMax may bring other mechanistic advantages. Based on our study of MafB, we know that BZIP factors can also hinder the propagation of torsional stress along DNA. Thus, the tandem binding of BHLH and BZIP factors could further enhance the transient accumulation of torsional stress, which could be necessary for the destabilisation of nucleosome core particles, the pre-initiation complex formation(Corless and Gilbert 2016), or DNA looping(Yan et al. 2018). The analyses of Myc (Figure S19A) and Max (Figure S19B) show both similar and different co-binding partners, which relates to the fact that Max forms both homodimers and heterodimers.

In summary, using atomistic microsecond range umbrella sampling simulations with the torsional restraint that controls DNA total twist, we have shown that BHLH TFs may hinder the propagation of torsional stress along DNA. When complexed with the homologous MycMax, MadMax, and MaxMax dimers, DNA show a similarly increase in the torsional rigidity but experience distinct torque-induced deformations, which may modulate the binding of collaborative TFs. We thus propose that changing torsional constraints on DNA may contribute to the differential transcriptional programs of homologous TFs.

## Supporting information

Supplementary movie DNA

Supplementary movie DNA-MadMax complex

Supplementary movie DNA-MaxMax complex

Supplementary movie DNA-MycMax complex

## Acknowledgements

The authors thank Swedish National Infrastructure for Computing (SNIC) for the generous provision of computing resources.

## Author Contributions

JH and AR conceived and designed the study. JH performed the MD simulations. KM performed the bioinformatics analyses. JH, KM, and AR analyzed the data. JH and AR wrote the article.

## Data availability statement

All data generated and analysed in this study are available from the corresponding author upon request.

## References

Abraham, Mark James, Teemu Murtola, Roland Schulz, Szilárd Páll, Jeremy C Smith, Berk Hess, and Erik Lindahl. 2015. “GROMACS: High Performance Molecular Simulations through Multi-Level Parallelism from Laptops to Supercomputers.” SoftwareX 1–2: 19–25. https://doi.org/10.1016/j.softx.2015.06.001.

Atchley, W R, and W M Fitch. 1997. “A Natural Classification of the Basic Helix-Loop-Helix Class of Transcription Factors.” Proceedings of the National Academy of Sciences of the United States of America 94 (10): 5172–76. https://doi.org/10.1073/pnas.94.10.5172.

Balaceanu, Alexandra, Diana Buitrago, Jürgen Walther, Adam Hospital, Pablo D Dans, and Modesto Orozco. 2019. “Modulation of the Helical Properties of DNA: Next-to-Nearest Neighbour Effects and Beyond.” Nucleic Acids Research 47 (24): 4418–4430.

Berendsen, H J C, J P M Postma, W F van Gunsteren, A DiNola, and J R Haak. 1984. “Molecular Dynamics with Coupling to an External Bath.” The Journal of Chemical Physics 81 (8): 3684–90. https://doi.org/10.1063/1.448118.

Boeger, Hinrich, David A Bushnell, Ralph Davis, Joachim Griesenbeck, Yahli Lorch, J Seth Strattan, Kenneth D Westover, and Roger D Kornberg. 2005. “Structural Basis of Eukaryotic Gene Transcription.” FEBS Letters 579 (4): 899–903.

Bonomi, Massimiliano, Davide Branduardi, Giovanni Bussi, Carlo Camilloni, Davide Provasi, Paolo Raiteri, Davide Donadio, et al. 2009. “PLUMED: A Portable Plugin for Free-Energy Calculations with Molecular Dynamics.” Computer Physics Communications 180 (10): 1961–72. https://doi.org/10.1016/j.cpc.2009.05.011.

Core Team, RCTR. 2013. “R: A Language and Environment for Statistical Computing.” R Foundation for Statistical Computing, Vienna.

Corless, Samuel, and Nick Gilbert. 2016. “Effects of DNA Supercoiling on Chromatin Architecture.” Biophysical Reviews 8 (3): 245–58. https://doi.org/10.1007/s12551-016-0210-1.

Dans, Pablo D, Alexandra Balaceanu, Marco Pasi, Alessandro S Patelli, Daiva Petkevičiūtė, Jürgen Walther, Adam Hospital, et al. 2019. “The Static and Dynamic Structural Heterogeneities of B-DNA: Extending Calladine–Dickerson Rules.” Nucleic Acids Research 47 (21): 11090–102. https://doi.org/10.1093/nar/gkz905.

Dans, Pablo D, Alberto Pérez, Ignacio Faustino, Richard Lavery, and Modesto Orozco. 2012. “Exploring Polymorphisms in B-DNA Helical Conformations.” Nucleic Acids Research 40 (21): 10668–78. https://doi.org/10.1093/nar/gks884.

Dennis, Daniel J, Sisu Han, and Carol Schuurmans. 2019. “BHLH Transcription Factors in Neural Development, Disease, and Reprogramming.” Brain Research 1705: 48–65. https://doi.org/10.1016/j.brainres.2018.03.013.

Diolaiti, Daniel, Lisa McFerrin, Patrick A Carroll, and Robert N Eisenman. 2015. “Functional Interactions among Members of the MAX and MLX Transcriptional Network during Oncogenesis.” Biochimica et Biophysica Acta 1849 (5): 484–500. https://doi.org/10.1016/j.bbagrm.2014.05.016.

Dunaway, Marietta, and Elaine A Ostrander. 1993. “Local Domains of Supercoiling Activate a Eukaryotic Promoter in Vivo.” Nature 361 (6414): 746–48.

Ferré-D’Amaré, Adrian R, George C Prendergast, Edward B Ziff, and Stephen K Burley. 1993. “Recognition by Max of Its Cognate DNA through a Dimeric b/HLH/Z Domain.” Nature 363 (6424): 38–45. https://doi.org/10.1038/363038a0.

Fogg, Jonathan M, Allison K Judge, Erik Stricker, Hilda L Chan, and Lynn Zechiedrich. 2021. “Supercoiling and Looping Promote DNA Base Accessibility and Coordination among Distant Sites.” Nature Communications 12 (1): 5683. https://doi.org/10.1038/s41467-021-25936-2.

Giardino Torchia, Maria Letizia, and Jonathan D Ashwell. 2018. “Getting MAD at MYC.” Proceedings of the National Academy of Sciences 115 (40): 9821 LP–9823. https://doi.org/10.1073/pnas.1813867115.

Grandori, Carla, Shaun M Cowley, Leonard P James, and Robert N Eisenman. 2000. “The Myc/Max/Mad Network and the Transcriptional Control of Cell Behavior.” Annual Review of Cell and Developmental Biology 16 (1): 653–99. https://doi.org/10.1146/annurev.cellbio.16.1.653.

Hörberg, J., and A. Reymer. 2020. “Specifically Bound BZIP Transcription Factors Modulate DNA Supercoiling Transitions.” Scientific Reports 10 (1). https://doi.org/10.1038/s41598-020-75711-4.

Hörberg, Johanna, Kevin Moreau, Markus J Tamás, and Anna Reymer. 2021. “Sequence-Specific Dynamics of DNA Response Elements and Their Flanking Sites Regulate the Recognition by AP-1 Transcription Factors.” Nucleic Acids Research, August. https://doi.org/10.1093/nar/gkab691.

Hörberg, Johanna, and Anna Reymer. 2018. “A Sequence Environment Modulates the Impact of Methylation on the Torsional Rigidity of DNA.” Chemical Communications 54 (84): 11885–88. https://doi.org/10.1039/C8CC06550K.

Ivani, Ivan, Pablo D Dans, Agnes Noy, Alberto Pérez, Ignacio Faustino, Adam Hospital, Jürgen Walther, et al. 2016. “Parmbsc1: A Refined Force Field for DNA Simulations.” Nature Methods 13 (1): 55–58. https://doi.org/10.1038/nmeth.3658.

Kannan, Srinivasaraghavan, Kai Kohlhoff, and Martin Zacharias. 2006. “B-DNA under Stress: Over- and Untwisting of DNA during Molecular Dynamics Simulations.” Biophysical Journal 91 (8): 2956–65. https://doi.org/10.1529/biophysj.106.087163.

Kouzine, Fedor, Ashutosh Gupta, Laura Baranello, Damian Wojtowicz, Khadija Ben-Aissa, Juhong Liu, Teresa M Przytycka, and David Levens. 2013. “Transcription-Dependent Dynamic Supercoiling Is a Short-Range Genomic Force.” Nature Structural & Molecular Biology 20 (3): 396–403. https://doi.org/10.1038/nsmb.2517.

Kumar, Shankar, John M Rosenberg, Djamal Bouzida, Robert H Swendsen, and Peter A Kollman. 1992. “THE Weighted Histogram Analysis Method for Free-Energy Calculations on Biomolecules. I. The Method.” Journal of Computational Chemistry 13 (8): 1011–21. https://doi.org/10.1002/jcc.540130812.

Lavelle, Christophe. 2008. “DNA Torsional Stress Propagates through Chromatin Fiber and Participates in Transcriptional Regulation.” Nature Structural & Molecular Biology 15 (2): 123–25. https://doi.org/10.1038/nsmb0208-123.

Lavery, R, M Moakher, J H Maddocks, D Petkeviciute, and K Zakrzewska. 2009. “Conformational Analysis of Nucleic Acids Revisited: Curves+.” Nucleic Acids Research 37 (17): 5917–29. https://doi.org/10.1093/nar/gkp608.

Lavery, R, K Zakrzewska, and H Sklenar. 1995. “JUMNA (Junction Minimisation of Nucleic Acids).” Computer Physics Communications 91 (1): 135–58. https://doi.org/10.1016/0010-4655(95)00046-I.

Liebl, Korbinian, and Martin Zacharias. 2017. “Unwinding Induced Melting of Double-Stranded DNA Studied by Free Energy Simulations.” The Journal of Physical Chemistry B 121 (49): 11019–30. https://doi.org/10.1021/acs.jpcb.7b07701.

Liebl, Korbinian, and Martin Zacharias. 2020. “How Global DNA Unwinding Causes Non-Uniform Stress Distribution and Melting of DNA.” PLOS ONE 15 (5): e0232976. https://doi.org/10.1371/journal.pone.0232976.

Liebl, Korbinian, and Martin Zacharias. 2021. “Accurate Modeling of DNA Conformational Flexibility by a Multivariate Ising Model.” Proceedings of the National Academy of Sciences 118 (15): e2021263118. https://doi.org/10.1073/pnas.2021263118.

Lin, Quy Xiao Xuan, Denis Thieffry, Sudhakar Jha, and Touati Benoukraf. 2020. “TFregulomeR Reveals Transcription Factors’ Context-Specific Features and Functions.” Nucleic Acids Research 48 (2): e10–e10. https://doi.org/10.1093/nar/gkz1088.

Lourenco, Corey, Diana Resetca, Cornelia Redel, Peter Lin, Alannah S MacDonald, Roberto Ciaccio, Tristan M G Kenney, et al. 2021. “MYC Protein Interactors in Gene Transcription and Cancer.” Nature Reviews Cancer 21 (9): 579–91. https://doi.org/10.1038/s41568-021-00367-9.

Ma, Jie, Lu Bai, and Michelle D Wang. 2013. “Transcription under Torsion.” Science (New York, N.Y.) 340 (6140): 1580–83. https://doi.org/10.1126/science.1235441.

Maier, James A, Carmenza Martinez, Koushik Kasavajhala, Lauren Wickstrom, Kevin E Hauser, and Carlos Simmerling. 2015. “Ff14SB: Improving the Accuracy of Protein Side Chain and Backbone Parameters from Ff99SB.” Journal of Chemical Theory and Computation 11 (8): 3696–3713. https://doi.org/10.1021/acs.jctc.5b00255.

Mark, Pekka, and Lennart Nilsson. 2001. “Structure and Dynamics of the TIP3P, SPC, and SPC/E Water Models at 298 K.” The Journal of Physical Chemistry A 105 (43): 9954–60. https://doi.org/10.1021/jp003020w.

Martin, Xabier de, Reza Sodaei, and Gabriel Santpere. 2021. “Mechanisms of Binding Specificity among BHLH Transcription Factors.” International Journal of Molecular Sciences. https://doi.org/10.3390/ijms22179150.

Masi, Federico De, Christian A Grove, Anastasia Vedenko, Andreu Alibés, Stephen S Gisselbrecht, Luis Serrano, Martha L Bulyk, and Albertha J M Walhout. 2011. “Using a Structural and Logics Systems Approach to Infer BHLH–DNA Binding Specificity Determinants.” Nucleic Acids Research 39 (11): 4553–63. https://doi.org/10.1093/nar/gkr070.

Mizutani, Mitsuko, Tsutomu Ohta, Hajime Watanabe, Hiroshi Handa, and Susumu Hirose. 1991. “Negative Supercoiling of DNA Facilitates an Interaction between Transcription Factor IID and the Fibroin Gene Promoter.” Proceedings of the National Academy of Sciences 88 (3): 718–22.

Mizutani, Mitsuko, Kiyoe Ura, and Susumu Hirose. 1991. “DNA Superhelicity Affects the Formation of Transcription Preinitiation Complex on Eukaryotic Genes Differently.” Nucleic Acids Research 19 (11): 2907–11.

Nair, Satish K, and Stephen K Burley. 2003. “X-Ray Structures of Myc-Max and Mad-Max Recognizing DNA: Molecular Bases of Regulation by Proto-Oncogenic Transcription Factors.” Cell 112 (2): 193–205. https://doi.org/10.1016/S0092-8674(02)01284-9.

Naughton, Catherine, Nicolaos Avlonitis, Samuel Corless, James G Prendergast, Ioulia K Mati, Paul P Eijk, Scott L Cockroft, Mark Bradley, Bauke Ylstra, and Nick Gilbert. 2013. “Transcription Forms and Remodels Supercoiling Domains Unfolding Large-Scale Chromatin Structures.” Nature Structural & Molecular Biology 20 (3): 387–95. https://doi.org/10.1038/nsmb.2509.

Noy, Agnes, Thana Sutthibutpong, and Sarah A. Harris. 2016. “Protein/DNA Interactions in Complex DNA Topologies: Expect the Unexpected.” Biophysical Reviews 8 (3): 233–43. https://doi.org/10.1007/s12551-016-0208-8.

Osborne, Cameron S, Lyubomira Chakalova, Karen E Brown, David Carter, Alice Horton, Emmanuel Debrand, Beatriz Goyenechea, Jennifer A Mitchell, Susana Lopes, and Wolf Reik. 2004. “Active Genes Dynamically Colocalize to Shared Sites of Ongoing Transcription.” Nature Genetics 36 (10): 1065–71.

Parrinello, M, and A Rahman. 1981. “Polymorphic Transitions in Single Crystals: A New Molecular Dynamics Method.” Journal of Applied Physics 52 (12): 7182–90. https://doi.org/10.1063/1.328693.

Pasi, Marco, John H Maddocks, David Beveridge, Thomas C Bishop, David a Case, Thomas Cheatham, Pablo D Dans, et al. 2014. “MABC: A Systematic Microsecond Molecular Dynamics Study of Tetranucleotide Sequence Effects in B-DNA.” Nucleic Acids Research 42 (19): 12272–83. https://doi.org/10.1093/nar/gku855.

Patel, Anamika, Peng Yang, Matthew Tinkham, Mihika Pradhan, Ming-An Sun, Yixuan Wang, Don Hoang, et al. 2018. “DNA Conformation Induces Adaptable Binding by Tandem Zinc Finger Proteins.” Cell 173 (1): 221–233.e12. https://doi.org/10.1016/j.cell.2018.02.058.

Pettersen, Eric F, Thomas D Goddard, Conrad C Huang, Gregory S Couch, Daniel M Greenblatt, Elaine C Meng, and Thomas E Ferrin. 2004. “UCSF Chimera—A Visualization System for Exploratory Research and Analysis.” Journal of Computational Chemistry 25 (13): 1605–12. https://doi.org/10.1002/jcc.20084.

Pyne, Alice L B, Agnes Noy, Kavit H S Main, Victor Velasco-Berrelleza, Michael M Piperakis, Lesley A Mitchenall, Fiorella M Cugliandolo, et al. 2021. “Base-Pair Resolution Analysis of the Effect of Supercoiling on DNA Flexibility and Major Groove Recognition by Triplex-Forming Oligonucleotides.” Nature Communications 12 (1): 1053. https://doi.org/10.1038/s41467-021-21243-y.

Reymer, Anna, Krystyna Zakrzewska, and Richard Lavery. 2018. “Sequence-Dependent Response of DNA to Torsional Stress: A Potential Biological Regulation Mechanism.” Nucleic Acids Research 46 (4): 1684–94. https://doi.org/10.1093/nar/gkx1270.

Roe, Daniel R, and Thomas E Cheatham. 2013. “PTRAJ and CPPTRAJ: Software for Processing and Analysis of Molecular Dynamics Trajectory Data.” Journal of Chemical Theory and Computation 9 (7): 3084–95. https://doi.org/10.1021/ct400341p.

Schultz, Michael C, Steven J Brill, QIDA Ju, Rolf Sternglanz, and Ronald H Reeder. 1992. “Topoisomerases and Yeast RRNA Transcription: Negative Supercoiling Stimulates Initiation and Topoisomerase Activity Is Required for Elongation.” Genes & Development 6 (7): 1332–41.

Tabuchi, Hisahiro, and Susumu Hirose. 1988. “DNA Supercoiling Facilitates Formation of the Transcription Initiation Complex on the Fibroin Gene Promoter.” Journal of Biological Chemistry 263 (30): 15282–87.

Teves, Sheila S, and Steven Henikoff. 2014. “Transcription-Generated Torsional Stress Destabilizes Nucleosomes.” Nature Structural & Molecular Biology 21 (1): 88–94. https://doi.org/10.1038/nsmb.2723.

Torrie, G M, and J P Valleau. 1977. “Nonphysical Sampling Distributions in Monte Carlo Free-Energy Estimation: Umbrella Sampling.” Journal of Computational Physics 23 (2): 187–99. https://doi.org/10.1016/0021-9991(77)90121-8.

Wei, Yong, Diana Resetca, Zhe Li, Isak Johansson-Åkhe, Alexandra Ahlner, Sara Helander, Amelie Wallenhammar, et al. 2019. “Multiple Direct Interactions of TBP with the MYC Oncoprotein.” Nature Structural & Molecular Biology 26 (11): 1035–43. https://doi.org/10.1038/s41594-019-0321-z.

Weintraub, Harold, Pei Feng Cheng, and Kathleen Conrad. 1986. “Expression of Transfected DNA Depends on DNA Topology.” Cell 46 (1): 115–22.

Yan, Yan, Fenfei Leng, Laura Finzi, and David Dunlap. 2018. “Protein-Mediated Looping of DNA under Tension Requires Supercoiling.” Nucleic Acids Research 46 (5): 2370–79. https://doi.org/10.1093/nar/gky021.

