## Supplementary Information for "Homologous BHLH transcription factors induce distinct deformations of torsionally-stressed DNA: a potential transcription regulation mechanism"

SUPORTING INFORMATION

```
Myc        KRRTHNVLERQRRNELKRSFFALRDQIPELENNEKAPKVVILKKATAYILSV 407
Max        KRAHHNALERKRRDHIKDSFHSLRDSVPSLQGE-KASRAQILDKATEYIQYM 75
Mad        SRSTHNEMEKNRRAHLRLCLEKLKGLVPLGPESSRHTTLSLLTKAKLHIKKL 109
           .*  ** :*::** .:: .:  *:. :*    . :     :* **. :*  :
```

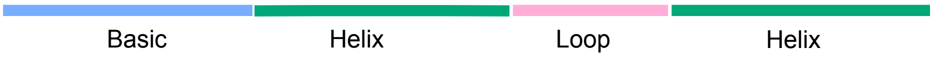


**Figure S1:** Multiple sequence alignment of the Basic-Helix-Loop-Helix domain of Myc, Max and Mad. The alignment has been performed using Clustal Omega.

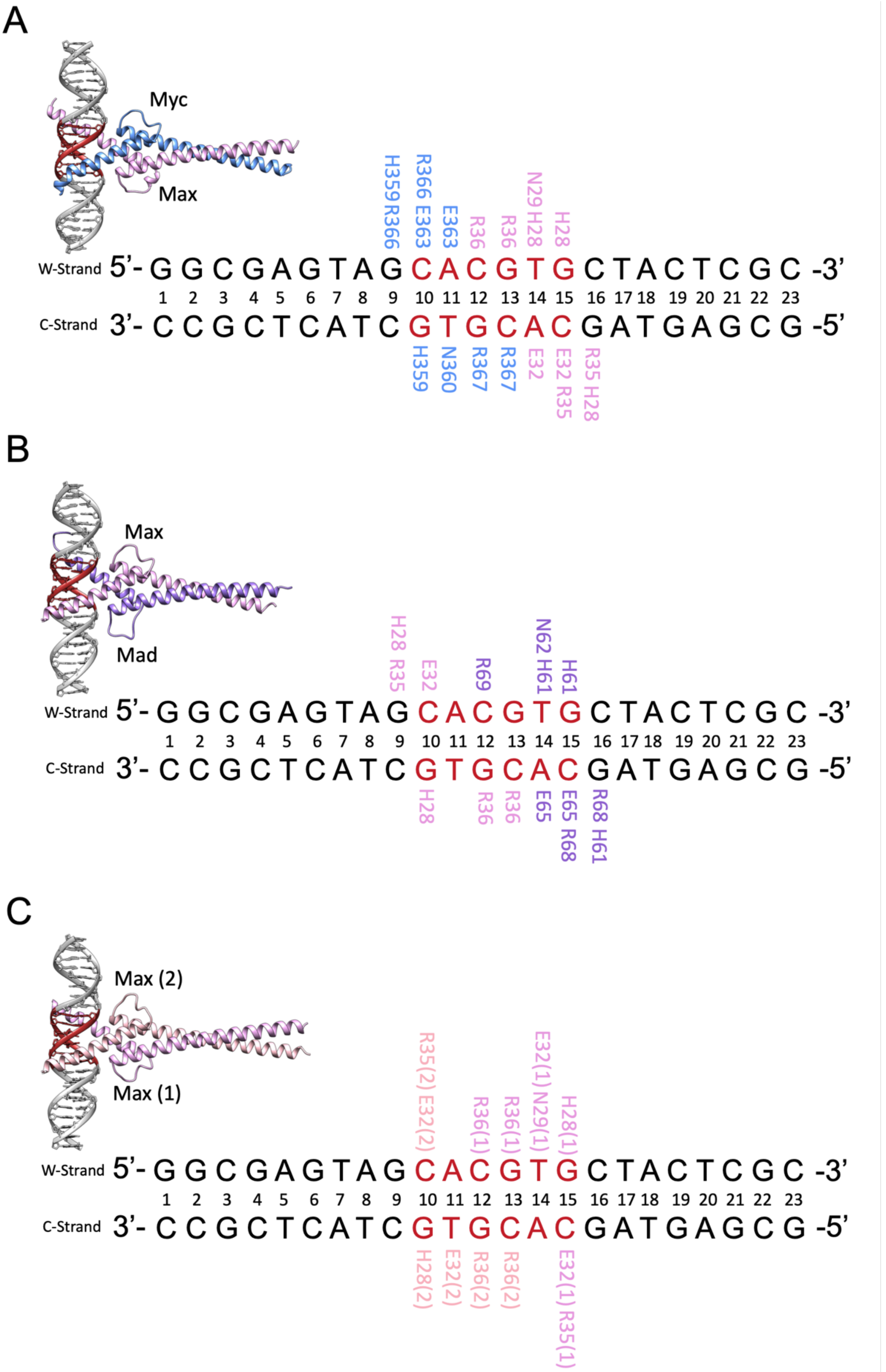


**Figure S2:** Crystal structure of **A.** MycMax-DNA (PDB ID: 1NKP), **B.** MadMax-DNA (PDB ID: 1NLW) and **C.** MaxMax-DNA (PDB ID: 1AN2) complexes bound to the E-Box element in red. For each complex, specific protein-DNA contacts seen in the crystal structures are highlighted. The Watson strand (5'->3') is denoted with "w" and the Crick (3'->5') strand is denoted with "c".

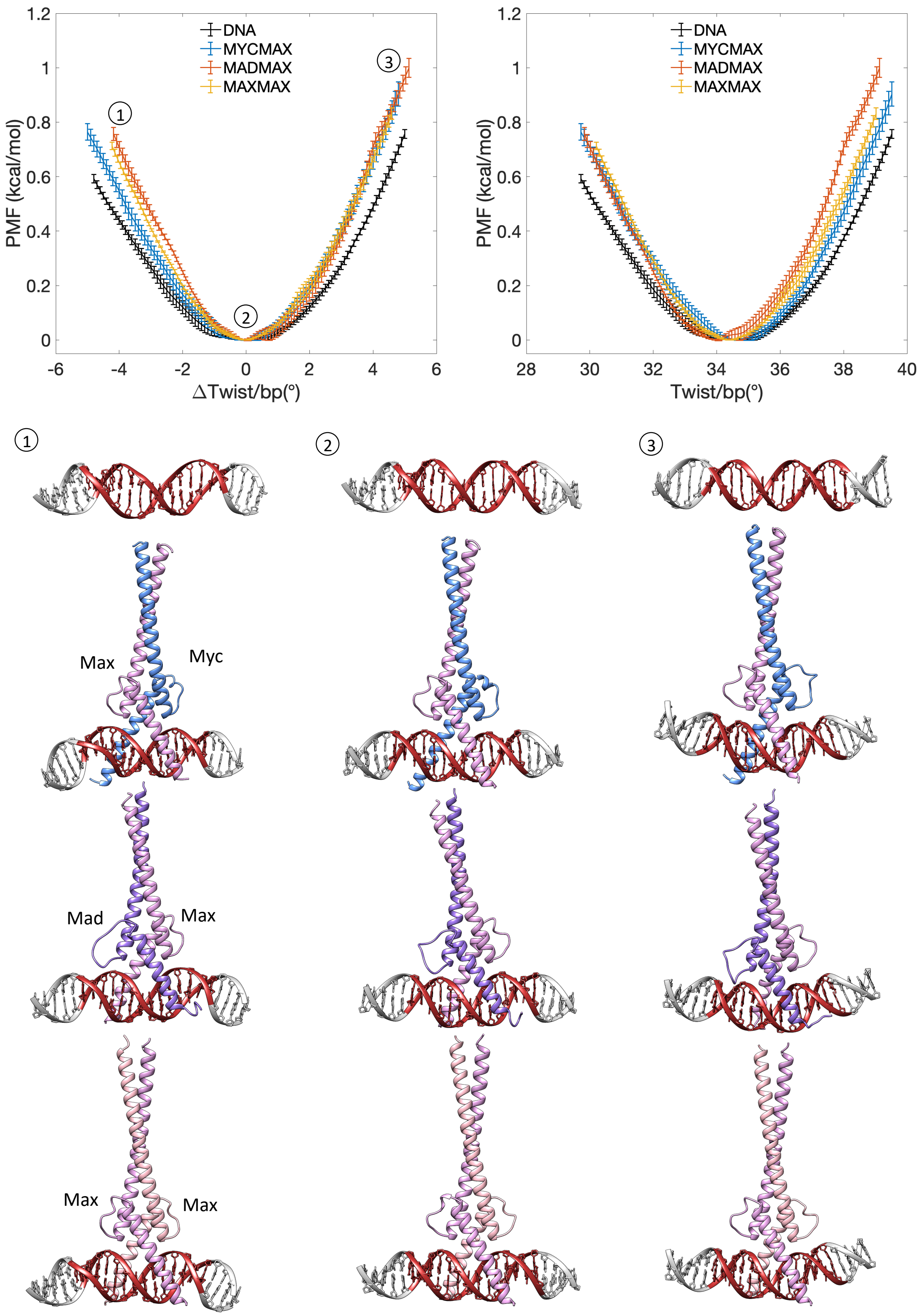


**Figure S3:** PMF profiles of DNA twisting transitions for unbound DNA and MycMax- (blue), MadMax- (orange) and MaxMax (yellow) bound DNA. Standard deviations of the 500 ns windows are highlighted as errorbars. DNA deformations for unbound DNA, MycMax-, MadMax- and MaxMax-bound DNA are shown for (1) underwound regime (-4.5°/bp), (2) torsionally relaxed regime and (3) overwound regime(+4.5°/bp). The restrained DNA region (GTAGCACGTGCTAC) is highlighted with red colour.

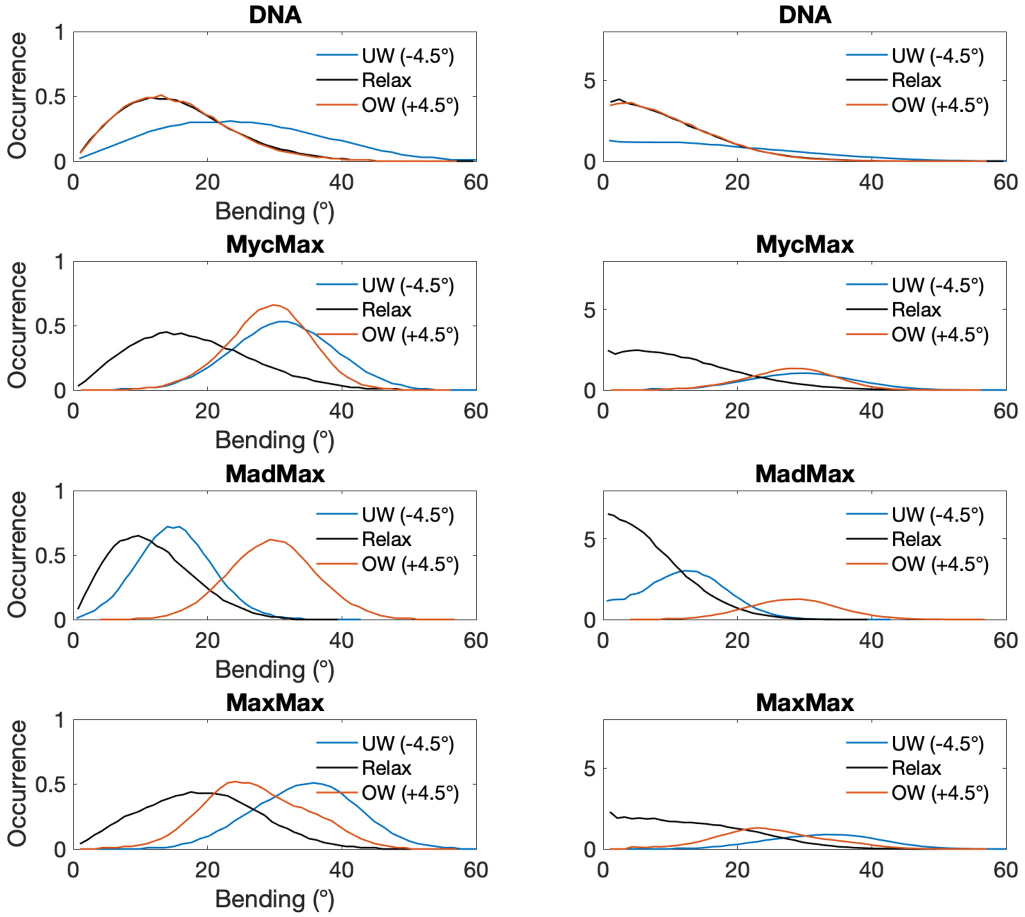


**Figure S4:** Bending distributions of the restrained region (GTAGCACGTGCTAC) for unbound DNA, MycMax-, MadMax- and MaxMax-bound DNA. For the right-hand panels, the histograms have been divided by sin($\boldsymbol{\theta}$), where $\boldsymbol{\theta}$ corresponds to the bending angle at the middle of each histogram bin, to compensate for the increasing area of the spherical ring segment sampled as the bending angle increases.

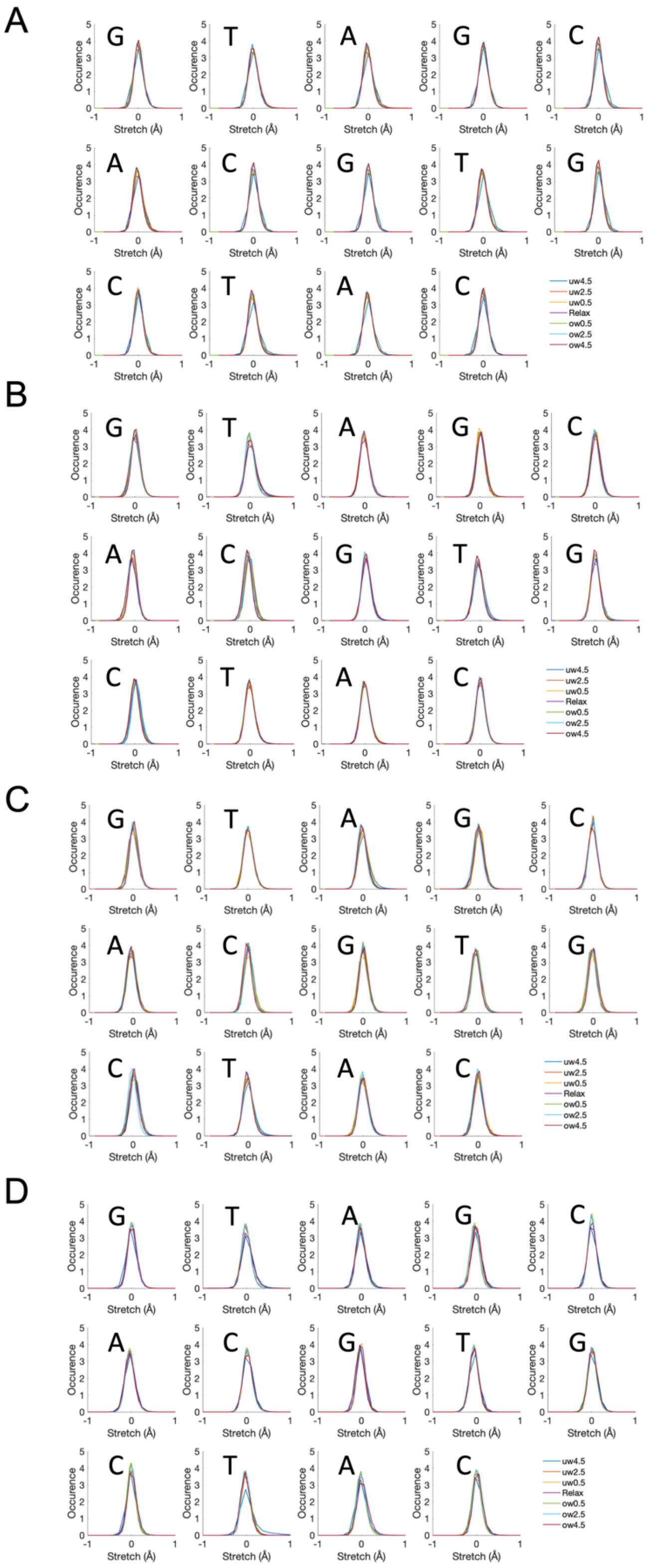


5’-GGCGA-**GTAGCACGTGCTAC**-TCGC-3’

**Figure S5:** Stretch distributions for b.p. of the restrained region for different torsional regimes. **A.** DNA, **B.** MycMax-DNA, **C.** MadMax-DNA, **D.** MaxMax-DNA.

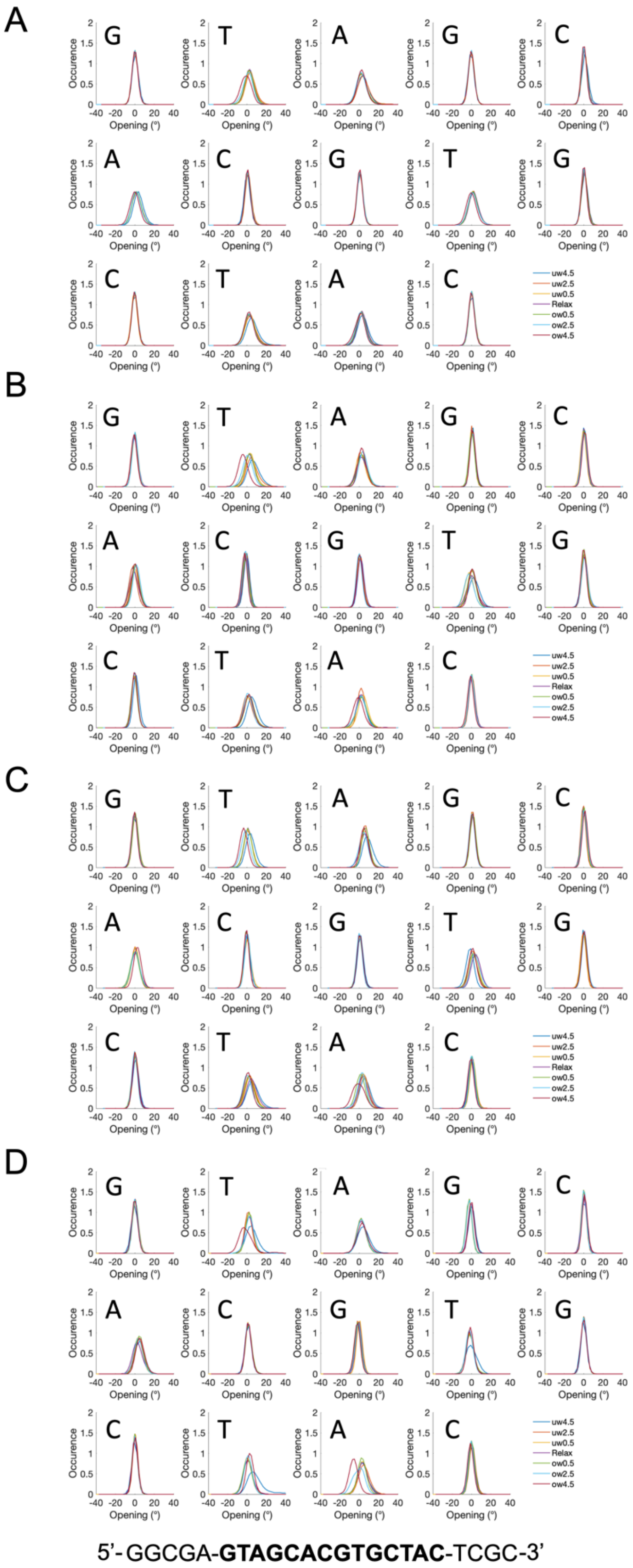


**Figure S6:** Opening distributions for b.p. of the restrained region for different torsional regimes. **A.** DNA, **B.** MycMax-DNA, **C.** MadMax-DNA, **D.** MaxMax-DNA.

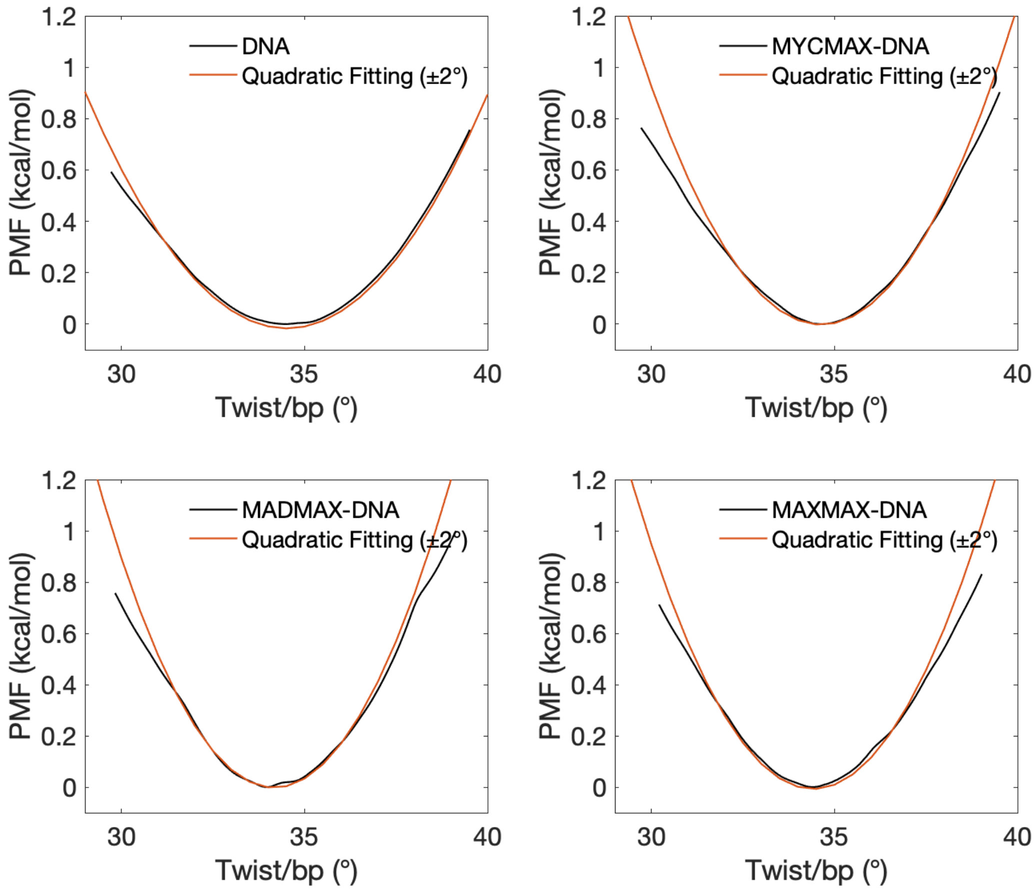


**Figure S7:** Quadratic regression (±2°) of PMF profiles of unbound DNA and MycMax-, MadMax- and MaxMax bound DNA is shown in orange lines.

**Table S1:** Calculated average relaxed twists, torsional constants ‘K’ (overall), ‘K-’ (undertwisting regime), ‘K+’ (overtwisting regime), torsional moduli ‘C’, and torsional persistence lengths ‘P’ for unbound DNA BHLH bound DNA.

| | DNA | MycMax-DNA | MadMax-DNA | MaxMax-DNA |
|---|---|---|---|---|
| Relaxed twist (°) | 34.5 | 34.7 | 34.0 | 34.4 |
| K (kcal/mol*deg$^2$) | 0.059 | 0.081 | 0.089 | 0.087 |
| K+ | 0.065 | 0.084 | 0.085 | 0.084 |
| K- | 0.042 | 0.051 | 0.056 | 0.054 |
| C (pN nm$^2$) | 457 | 628 | 691 | 672 |
| P (nm) | 111 | 153 | 168 | 163 |

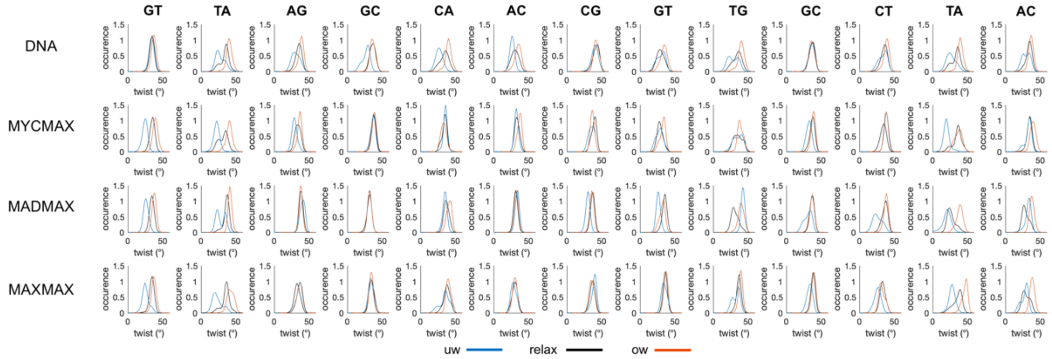


**Figure S8:** Twist distributions of the restrained DNA region (GTAGCACGTGCTAC) for underwound (-4.5), relaxed and overwound (+4.5) state.

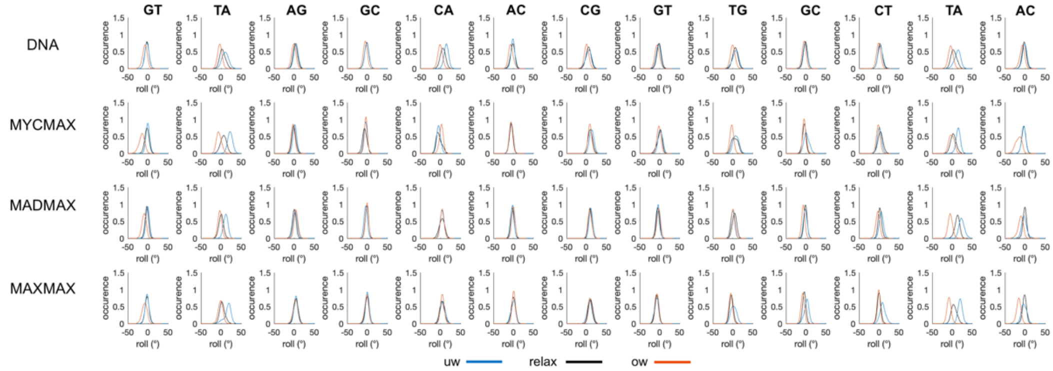


**Figure S9:** Roll distributions of the restrained DNA region (GTAGCACGTGCTAC) for underwound (-4.5), relaxed and overwound (+4.5) state.

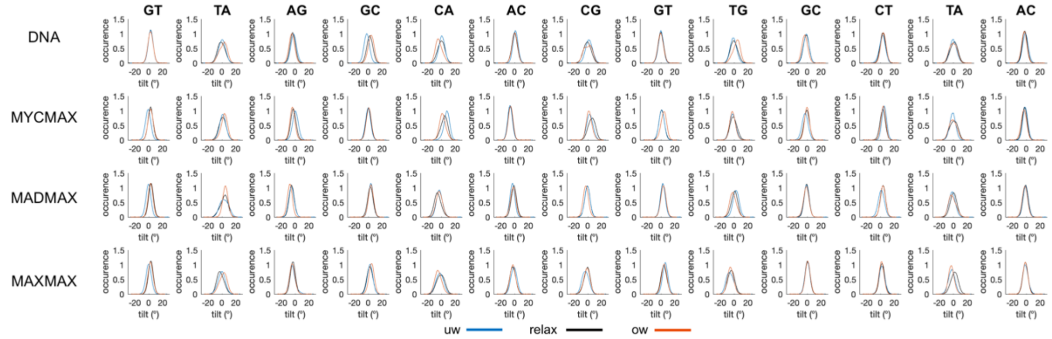


**Figure S10:** Tilt distributions of the restrained DNA region (GTAGCACGTGCTAC) for underwound (-4.5), relaxed and overwound (+4.5) state.

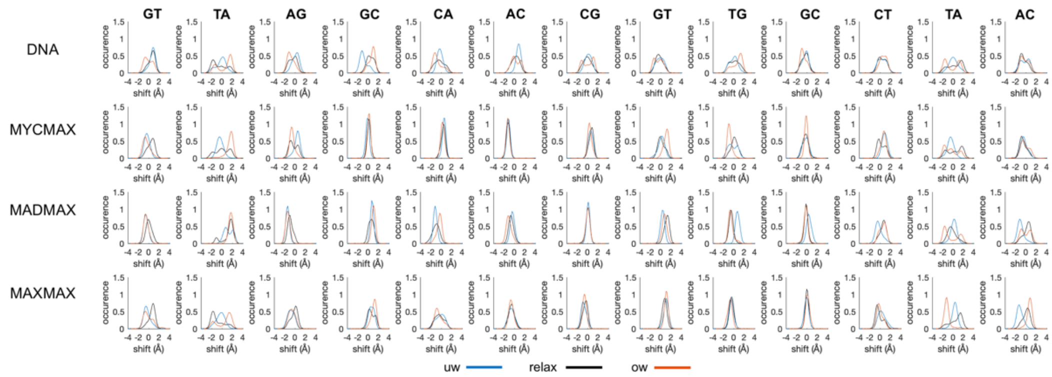


**Figure S11:** Shift distributions of the restrained DNA region (GTAGCACGTGCTAC) for underwound (-4.5), relaxed and overwound (+4.5) state.

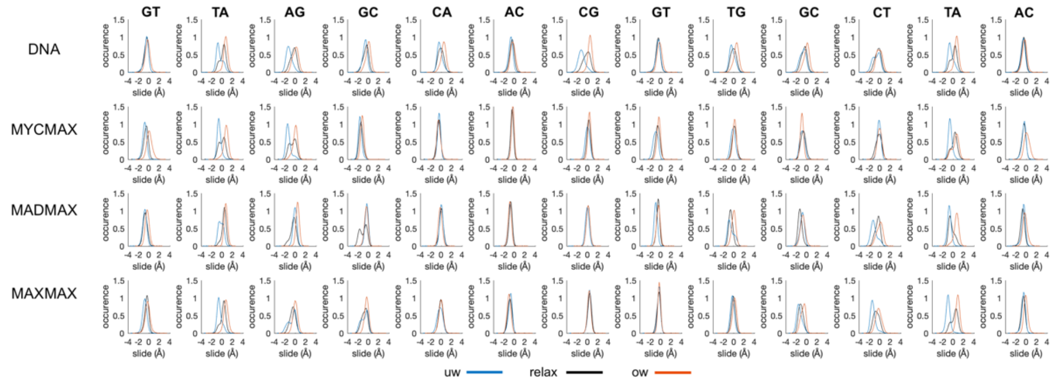


**Figure S12:** Slide distributions of the restrained DNA region (GTAGCACGTGCTAC) for underwound (-4.5), relaxed and overwound (+4.5) state.

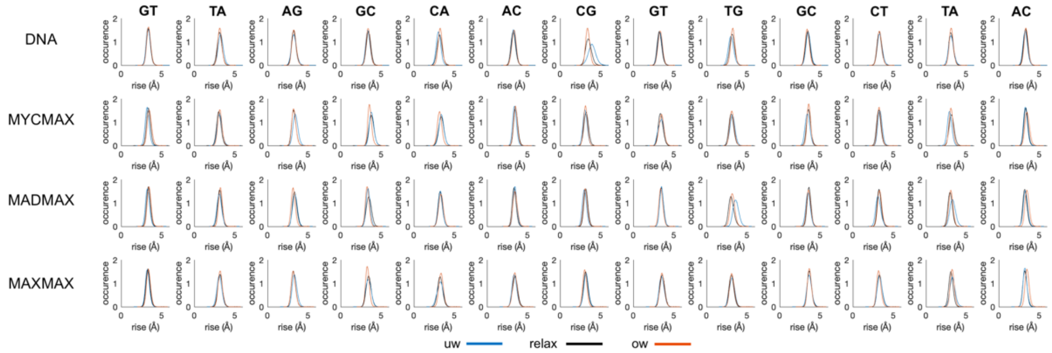


**Figure S13:** Rise distributions of the restrained DNA region (GTAGCACGTGCTAC) for underwound (-4.5), relaxed and overwound (+4.5) state.

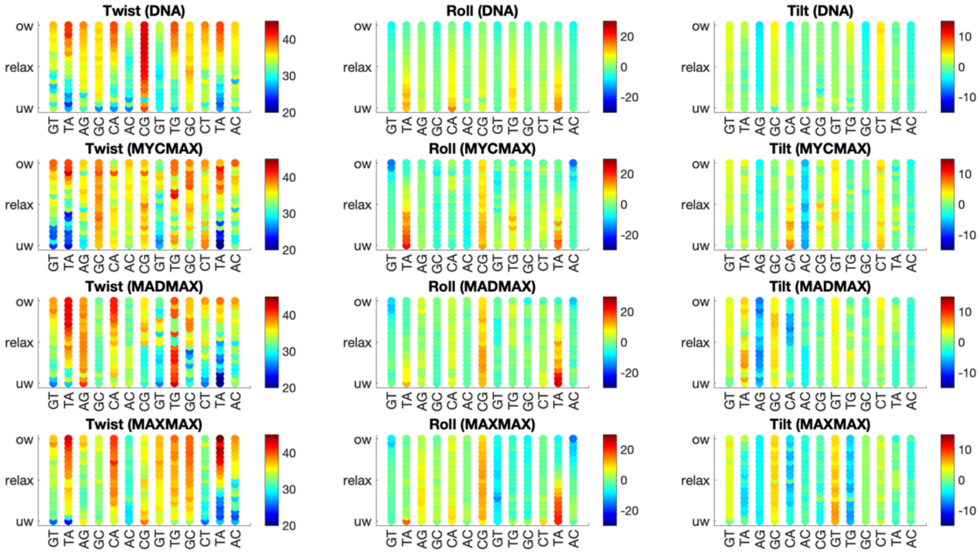


**Figure S14:** Change in average twist, roll and tilt of b.p. steps of the restrained DNA region (GTAGCACGTGCTAC) along the torsional regimes denoted with a colourbar.

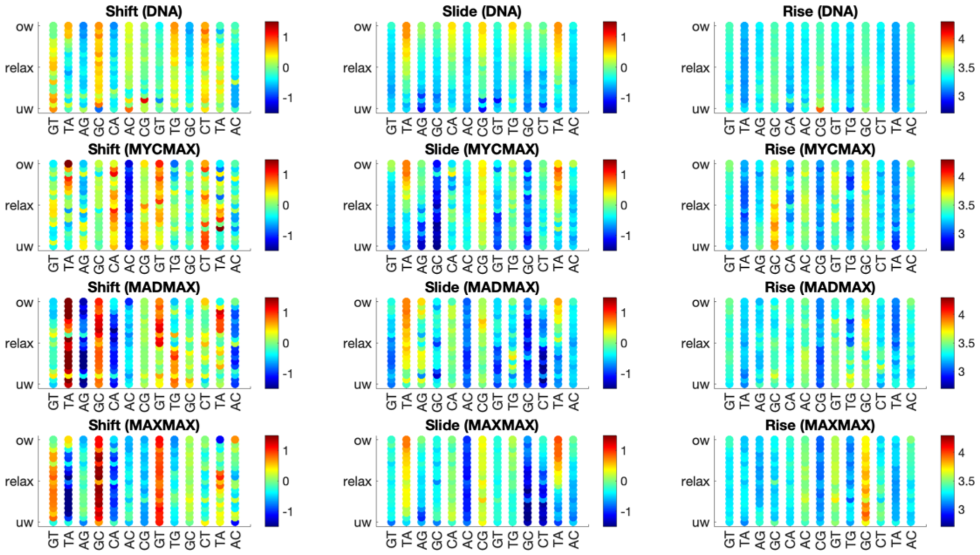


**Figure S15:** Change in average shift, slide and rise of b.p. steps of the restrained DNA region (GTAGCACGTGCTAC) along the torsional regimes denoted with a colourbar.

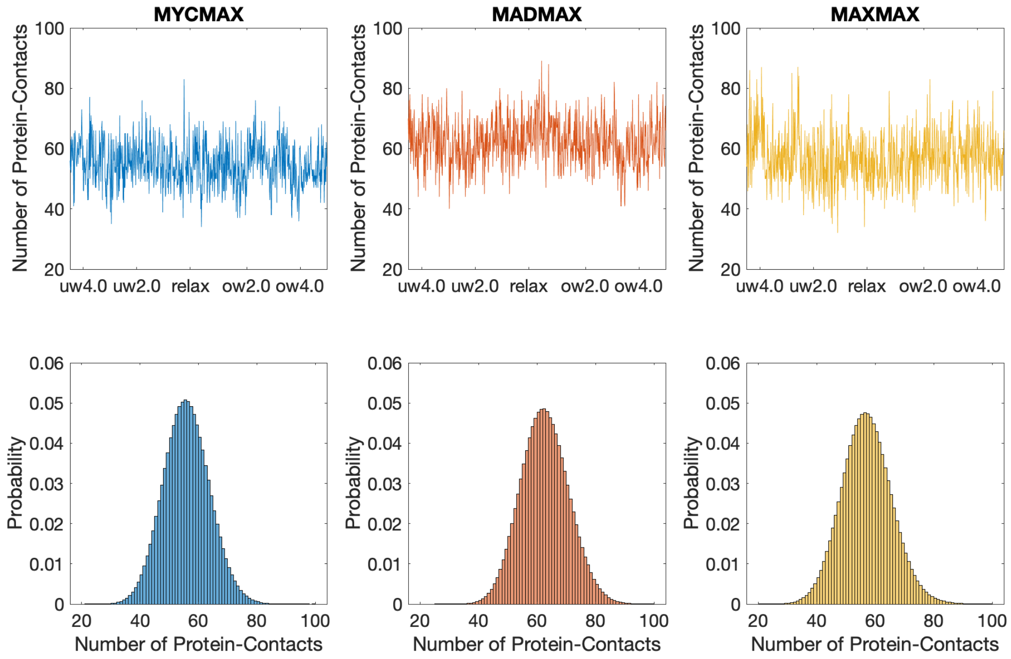


**Figure S16:** Upper panel: fluctuations of the total number of protein-DNA contacts along the torsional regimes. Bottom panel: distributions of the total number of protein-DNA contacts for the different BHLH-DNA complexes.

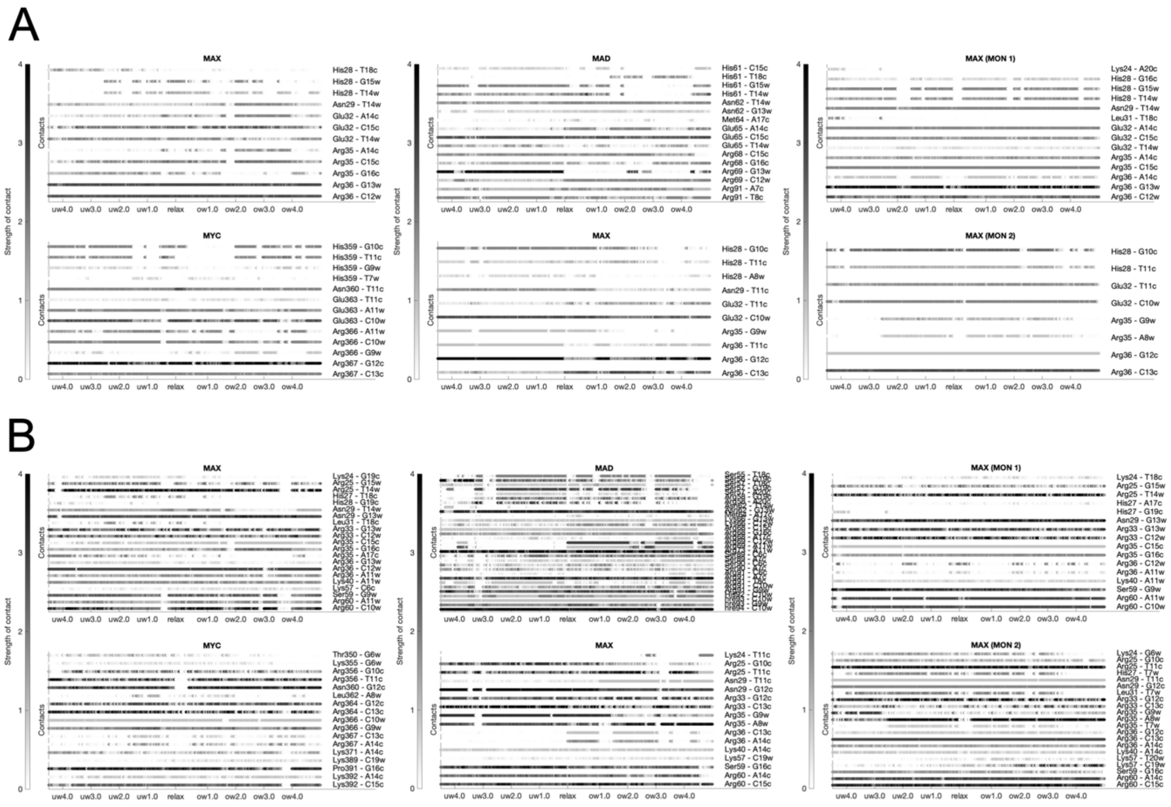


W-Strand 5'- G G C G A G T A G C A C G T G C T A C T C G C -3'
1 2 3 4 5 6 7 8 9 10 11 12 13 14 15 16 17 18 19 20 21 22 23
C-Strand 3'- C C G C T C A T C G T G C A C G A T G A G C G -5'

**Figure S17:** Dynamic interactions maps along the different torsional regimes for MycMax, MadMax and MaxMax. **A.** Strength of specific protein-DNA contacts. **B.** Strength of nonspecific protein-DNA contacts.

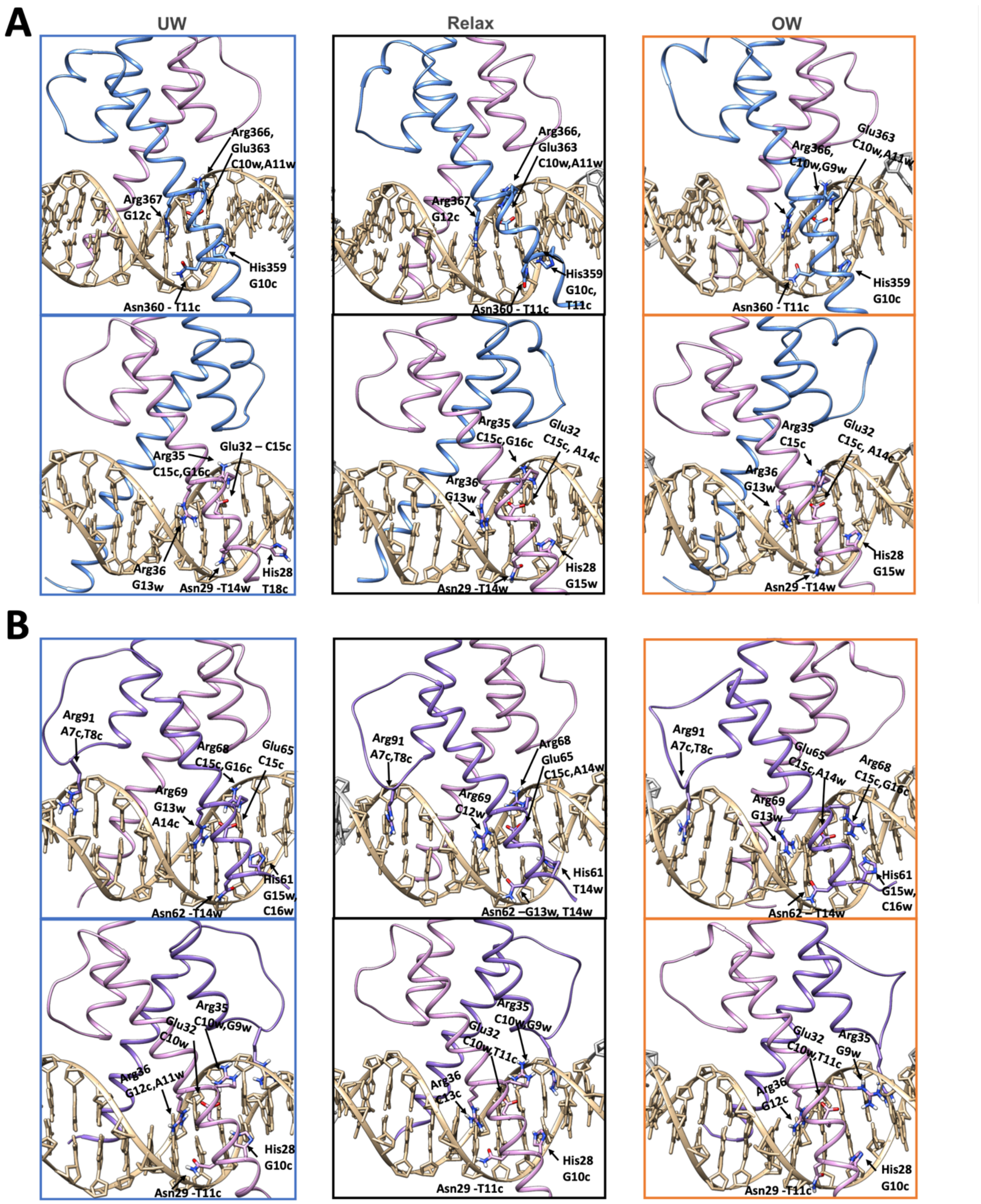

A
UW
Relax
OW
Arg366, Glu363 C10w,A11w
Arg367 G12c
His359 G10c
Asn360 - T11c
His359 G10c, T11c
Glu363 C10w,A11w
Arg366, C10w,G9w
Arg35 C15c,G16c
Glu32 – C15c
Arg36 G13w
Asn29 -T14w
His28 T18c
Arg35 C15c,G16c
Glu32 C15c, A14c
His28 G15w
Arg35 C15c
B
Arg91 A7c,T8c
Arg68 C15c,G16c
Glu65 C15c
Arg69 G13w A14c
His61 G15w, C16w
Asn62 -T14w
Arg68 Glu65 C15c,A14w
Arg69 C12w
His61 T14w
Asn62 –G13w, T14w
Glu65 C15c,A14w
Arg68 C15c,G16c
Arg69 G13w
Asn62 – T14w
Arg35 C10w,G9w
Glu32 C10w
Arg36 G12c,A11w
His28 G10c
Asn29 -T11c
Glu32 C10w,T11c
Arg36 C13c
Arg35 G9w
Arg36 G12c

**C**

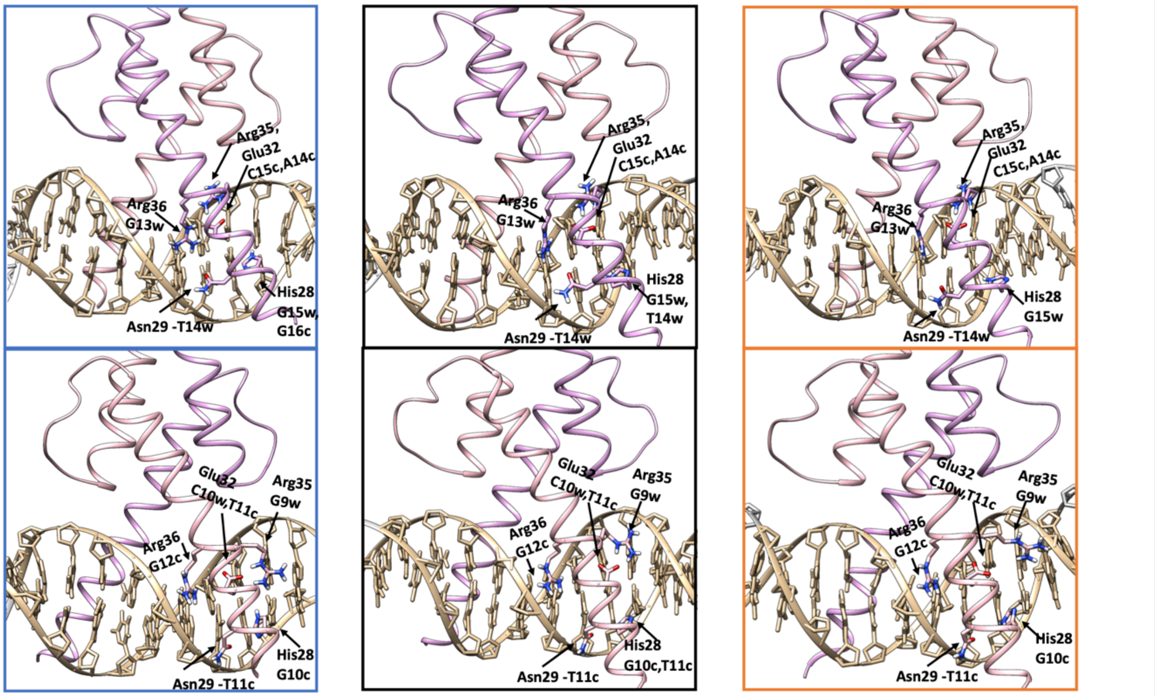


**Figure S18:** BHLH-DNA contacts exploited by the five-residues motif (**HN**xx**E**xx**RR**) for underwinding regime (-4.5°/bp), relaxed state and overwinding regime (+4.5°/bp). Restrained DNA region (GTAGCACGTGCTAC) is denoted with tan colour. **A.** Upper panel: Myc-DNA contacts. Bottom panel: Max-DNA contacts. **B.** Upper panel: Mad-DNA contacts. Specific contacts exploited by Mad loop is also shown. Bottom panel: Max-DNA contacts. **C.** Upper panel: Max(monomer 1)-DNA contacts. Bottom panel: Max(monomer 2)-DNA contacts.

A

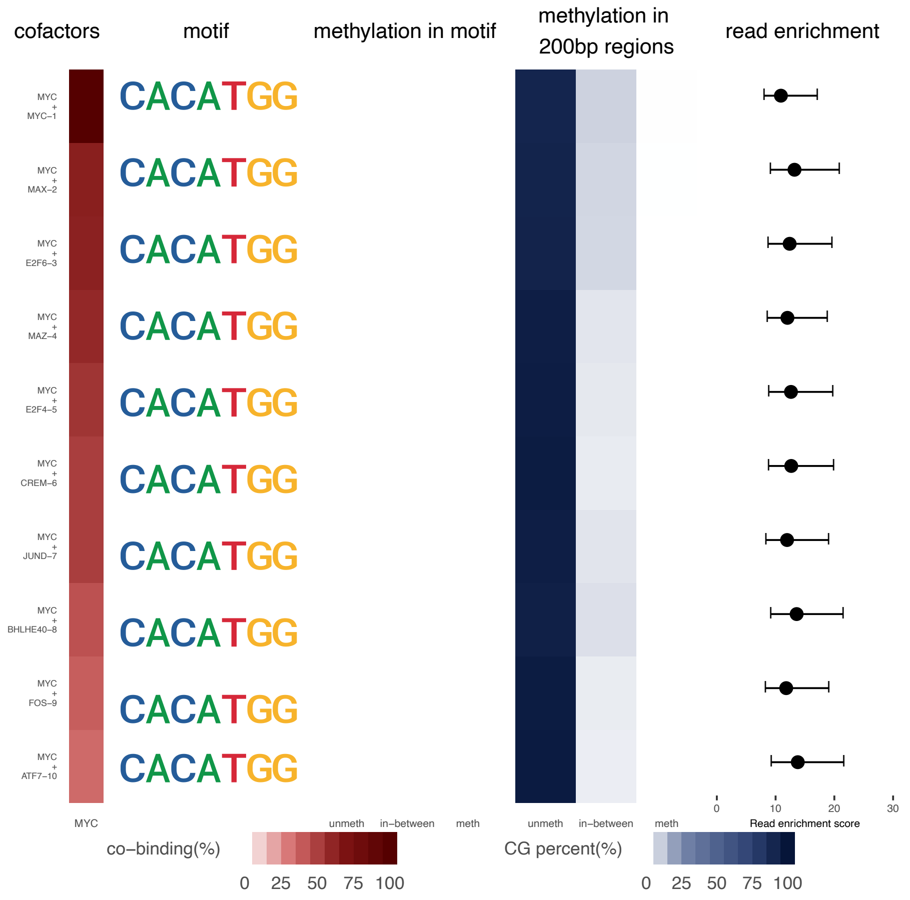

cofactors
motif
methylation in motif
methylation in 200bp regions
read enrichment
MYC + MYC−1
MYC + MAX−2
MYC + E2F6−3
MYC + MAZ−4
MYC + E2F4−5
MYC + CREM−6
MYC + JUND−7
MYC + BHLHE40−8
MYC + FOS−9
MYC + ATF7−10
CACATGG
MYC
unmeth
in−between
meth
Read enrichment score
0
10
20
30
co−binding(%)
0 25 50 75 100
CG percent(%)
0 25 50 75 100


B

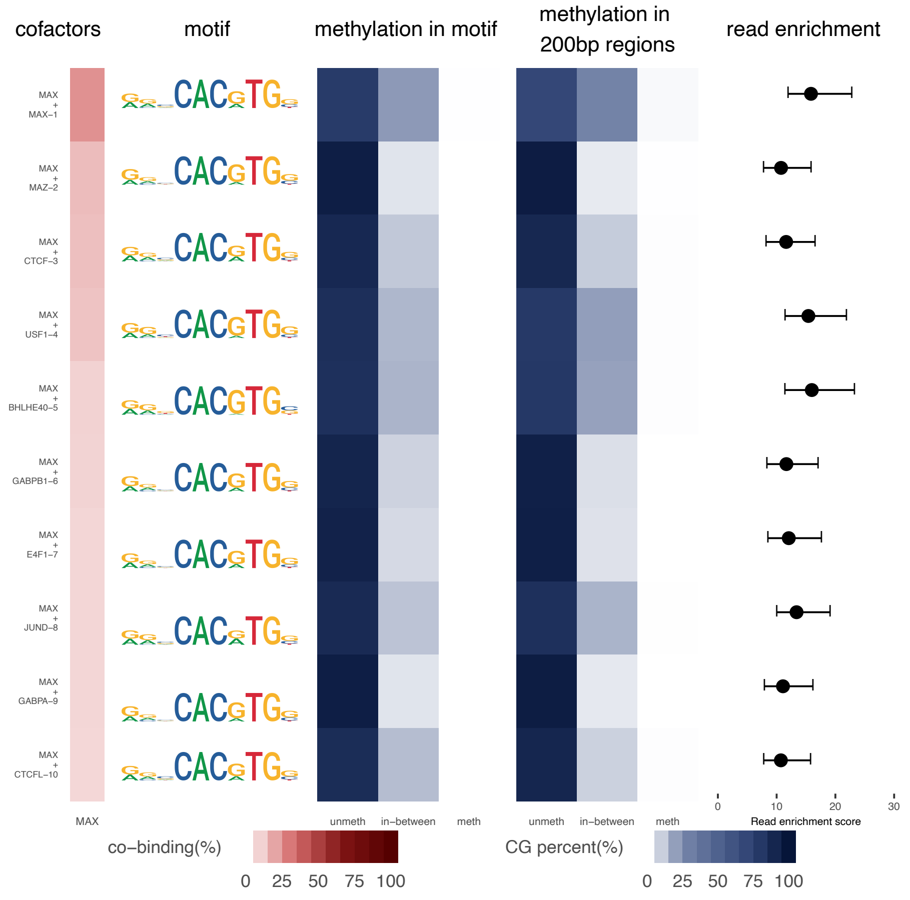

cofactors
motif
methylation in motif
methylation in 200bp regions
read enrichment
MAX + MAX−1
MAX + MAZ−2
MAX + CTCF−3
MAX + USF1−4
MAX + BHLHE40−5
MAX + GABPB1−6
MAX + E4F1−7
MAX + JUND−8
MAX + GABPA−9
MAX + CTCFL−10
MAX
unmeth
in−between
meth
Read enrichment score
0
10
20
30
co−binding(%)
0 25 50 75 100
CG percent(%)
0 25 50 75 100

**Figure S19: A.** Myc and **B.** Max co-factors summary as reported by TFregulomeR. Information reported are, from left to right, the name of transcription factors detected as potential co-factors to either Myc or Max, with the corresponding co-binding percentage in shade of red; the sequence motif for the main protein of interest (Myc or Max); the methylation percentage of cytosine inside the binding motif and in a window of 200bp around the binding motif, represented in shade of blue; the reads enrichment from the ChIP-seq profile of the considered proteins.
